# Breaking Free: Development of Circular AAV Cargos for Targeted Seamless Integration in the Liver

**DOI:** 10.1101/2024.10.26.620313

**Authors:** Brett J.G. Estes, Nisha Gandhi, Jessica Von Stetina, Dev Paudel, Angela X. Nan, Parth Amin, Joshua Rose, Shuai Wu, Kangni Zheng, Yijun Zhang, Jesse C. Cochrane, Jonathan D. Finn, Jenny Xie

## Abstract

Recent advancements in gene insertion have shifted from DNA repair-dependent mechanisms to more precise approaches, enhancing safety and predictability for editing outcomes. Integrase-mediated programmable genomic integration (I-PGI) utilizes a DNA cargo to insert transgenes in a targeted, unidirectional manner. *In vivo*, where nuclear delivery of DNA is challenging, adeno-associated virus (AAV) can act as the cargo vector. While I-PGI does not require DNA double-stranded breaks (DSBs) for activity, linear DNA cargo, like AAV, stimulates DNA end joining activity after integration. To mitigate potential risks from DSBs with linear viral cargo, we developed two circular genome types capable of seamless gene insertion in non-dividing cells. We first harnessed the orthogonal property of large serine integrases to produce circle-AAV (cAAV) from linear viral genomes in cells. cAAV demonstrated faithful seamless cargo integration in primary human hepatocytes (PHH) and robust DSB-free insertion structures *in vivo*. We then investigated the delivery of packaged circular AAV cargo (AAV.AD), which eliminates the need for enzymatic manipulation in the cell. AAV.AD proved to be a viable cargo for I-PGI, exhibiting functional integration in PHH and *in vivo*, that resulted in seamless insertion structures. Together, these findings provide the first reported evidence of DSB-free programmable genomic integration using integrase and AAV cargo, addressing a previously unrecognized challenge in the field.

## Introduction

Repurposing naturally occurring DNA-modifying enzymes has resulted in a diverse toolbox of gene insertion technologies for therapeutic advancements [1]. Targeted gene insertion at native genomic locations ensures stable, regulated expression and reduces the risks associated with non-native expression (too much or too little expression) [2, 3]. While programmable nucleases have facilitated various site-specific insertion strategies, they remain dependent on cellular DNA double-stranded break (DSB) repair and are significantly influenced by cell cycle state [4–9]. Although these strategies have shown efficacy, DSB-dependent gene insertion carries risk and can lead to heterologous insertion outcomes [10–12].

Large serine integrases (LSIs), like Bxb1, offer another avenue for seamless insertion due to their lack of DNA repair dependency [13–15]. LSIs mediate recombination between bacteria-derived (attB) and phage-derived (attP) attachment sites, resulting in a unidirectional insertion product flanked by attL and attR scars [13]. Recent efforts to reprogram Bxb1 to target the human genome have shown potential, but efficiencies in primary cells remain limited [16]. We recently developed integrase-mediated programmable genomic integration (I-PGI), derived from PASTE [17], which enables high-efficiency targeted gene insertion *in vivo* [18]. With I-PGI, the integrase beacon (attB or attP) is written or ligated [19] into the desired genomic location, allowing an engineered Bxb1 [20] to facilitate programmable genomic integration with a DNA cargo containing the corresponding attachment site. This process is specific to the central dinucleotide present in both attachment sites, which can also enable multiplexing capabilities [17, 21]. Aside from requiring double-stranded DNA substrates, Bxb1 has no specific cargo limitations and can integrate DNA sequences of any size, whether linear or circular, viral or non-viral [17, 22, 23]. While linear cargo substrates do not reduce recombination efficiency compared to circular, their insertion products result in a DSB repair dependency due to the lack of closed ends [24, 25].

The choice of I-PGI cargo format is driven by delivery feasibility into the target cell type. For hepatocytes, nuclear delivery of non-viral DNA remains challenging, making viral vectors like adeno-associated virus (AAV) the preferred approach for delivery [26, 27]. AAV is a linear 4,700 nucleotide single-stranded DNA genome (ssAAV) flanked by two inverted terminal repeats (ITRs) that are essential for genome replication and encapsidation [28–31]. During replication, linear genomes are primarily generated, although circular AAV intermediates have also been identified [32]. Moreover, replacement of both ITRs with a singular truncated ITR (AD domain) can facilitate single-stranded circular genome production through an alternative replication pathway [33]. Alternatively, mutation of one ITR can drive self-complimentary genome (scAAV) formation during production, leading to a double-stranded DNA genome that demonstrates stronger potency *in vivo* due to lack of second-strand synthesis requirements [34, 35]. While ssAAV and scAAV are initially linear at the time of transduction, ITR-driven recombination can convert genomes to circular episomes and heterologous concatemers over time [36, 37]. This process facilitates vector persistence *in vivo,* but is also very inefficient, with the vast majority of viral genomes being lost over time [38–40].

Despite the prevalence of AAV in genome editing, LSI-based recombination of AAV cargo has yet to be characterized. In this study, we demonstrate that Bxb1-mediated insertion of wildtype scAAV is DSB-dependent. To address this observation, we designed circle-AAV (cAAV) and AAV.AD cargo types that are capable of seamless DSB-free integration in the liver. To construct cAAV, we engineer AAV genomes to include additional Bxb1 attachments sites orthogonal to those used for targeted insertion. This approach leverages integrase multiplexing activity during I-PGI to drive both cargo sequence circularization and genomic integration of cAAV. We validate the production of circular dsDNA episomes from cAAV *in vitro,* demonstrate robust integration in both PHH and mice, and find that cAAV predominantly produces seamless integration products *in vivo*.

To produce AAV.AD, we take advantage of the ability of AAV to replicate circular products through a single ITR AD domain and investigate the production and delivery of circular viral cargo for I-PGI. We confirm functional viral production and determine packaged genomes exist in a circular state. We also show that AAV.AD are functional viral particles that result in faithful integrase-mediated insertion in both PHH and *in vivo*. We resolve the structure of integrated AAV.AD in PHH which confirms seamless integration products with this cargo type. Together, we demonstrate that circular viral cargos like cAAV and AAV.AD advance integrase-mediated targeted gene insertion by providing innovative, DSB-free solutions in tissues with cargo delivery challenges.

## Results

### DSB-dependent Integration with WT AAV

We integrated self-complimentary AAV (scAAV) into the PAH locus in primary human hepatocytes (PHH) with I-PGI. This scAAV cargo contains wildtype (WT) coding sequence, a splice site acceptor and a polyadenylation signal to facilitate transgene expression from the endogenous locus. This cargo sequence is flanked by inverted terminal repeats (ITRs), which are expected to integrate downstream of cargo sequence (Fig. 1a). To understand the DNA structure of integrase-mediated WT AAV insertion, we sequenced amplified insertion products with Oxford Nanopore Technologies (ONT) long-read sequencing. Read alignment revealed unidirectional gene insertion, strong alignment at attL and a deletion pattern within the expected ITR boundary region flanking attR (Fig. 1b). ONT targeted native genomic DNA sequencing reproduced the ITR deletion pattern, which enabled use of amplified DNA for greater sequencing depth in subsequent experiments (Extended Data Fig. 1). This suggests that linear AAV integration creates ITR-containing free ends at the attR junction, which is likely recognized as a DNA double-strand break (DSB), requiring DNA end joining to complete gene insertion (Fig. 1c).

**Figure 1.**
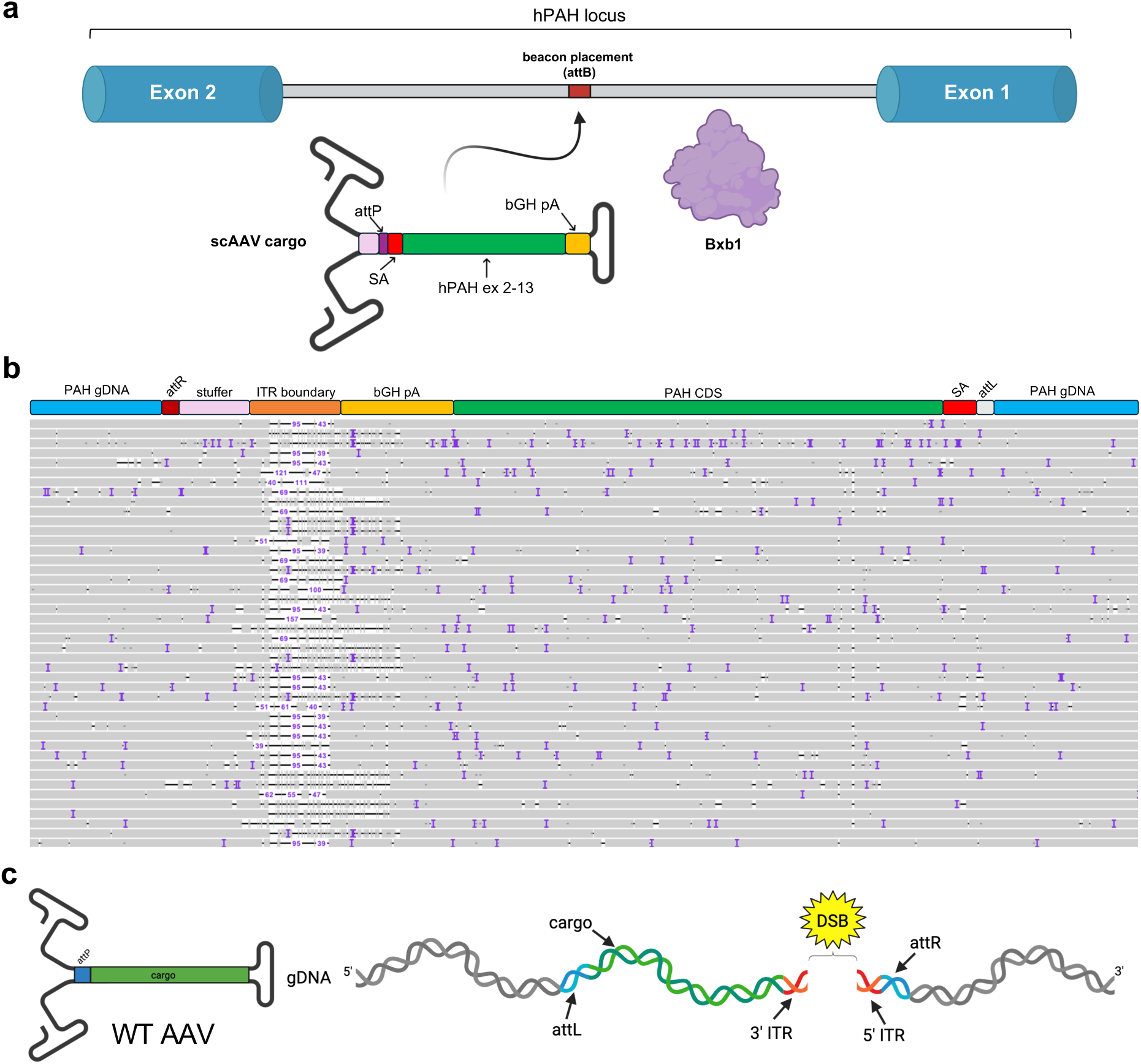
Integrase-mediated programmable genomic integration with wildtype AAV cargo. (**a**) An overview of I-PGI strategy for PKU targeting *PAH* intron 1. Beacons (attB) are placed within the intronic region where Bxb1 facilitates recombination with attP scAAV cargo. AAV cargo sequence includes a splice acceptor (SA), transgene coding sequence and polyadenylation signal (pA) that facilitate expression from the native locus. AAV genomes are flanked by inverted terminal repeats (ITRs) that integrate between the pA and attR sequences. (**b**) Representative long-read sequencing alignment of Bxb1-mediated insertion of WT scAAV cargo at the human *PAH* locus in PHH. Each line represents an individual read with alignment (grey) and deletions (black line) of various sizes (purple number) within the ITR boundary region. Samples were generated with a single I-PGI transfection of synthetic atgRNA, nCas9-RT mRNA and Bxb1 mRNA. (**c**) Schematic representing DNA repair dependency of WT AAV insertion products. Integration of WT linear AAV results in ITR-containing free ends at the attR junction that are likely recognized as a DNA double-stranded break (DSB).

### Development of cAAV

We developed circle-AAV (cAAV) with orthogonal Bxb1 attachment sites (attB*, attP*) flanking each ITR region to facilitate cargo circularization and DNA repair-independent integration (Fig. 2a). Integrases are capable of excision and intramolecular recombination within the cargo sequence would produce circular double-stranded DNA (dsDNA) episomes to enable seamless genomic insertion. Further, use of a distinct dinucleotide pair for circularization would ensure no cross talk between attachment sites utilized for genomic integration. To confirm that cAAV can circularize, we used an on/off dual luciferase single-stranded cAAV reporter where firefly luciferase is expressed in the linear, native format and NanoLuc is expressed upon circularization with Bxb1 (Fig. 2b). In PHH, we observed robust cAAV circularization in a Bxb1-dependent manner, as evidenced by reporter conversion from Firefly luciferase to Nanoluc expression (Fig. 2c). We also tested circularization of self-complimentary cAAV, which exhibited enhanced efficiencies compared to single-stranded cAAV (Extended Data Fig. 2). This suggests that the use of a more potent genome configuration could increase the concentration of circular episomes. cAAV circularization frequency (scAAV genome type) was assessed in PHH using droplet digital PCR (ddPCR) comparing circularized genomes to total genomes. A Bxb1 dose-response with cAAV showed a maximum circularization frequency of 10-15% (Fig. 2d). This frequency is likely an underestimate of the true efficiency given that there may have been AAV genomes that had yet to uncoat, making them inaccessible for integrase activity. Notably, cAAV functioned with alternative tyrosine/serine recombinases, further highlighting the modularity of this cargo system (Extended Data Fig. 3).

**Figure 2.**
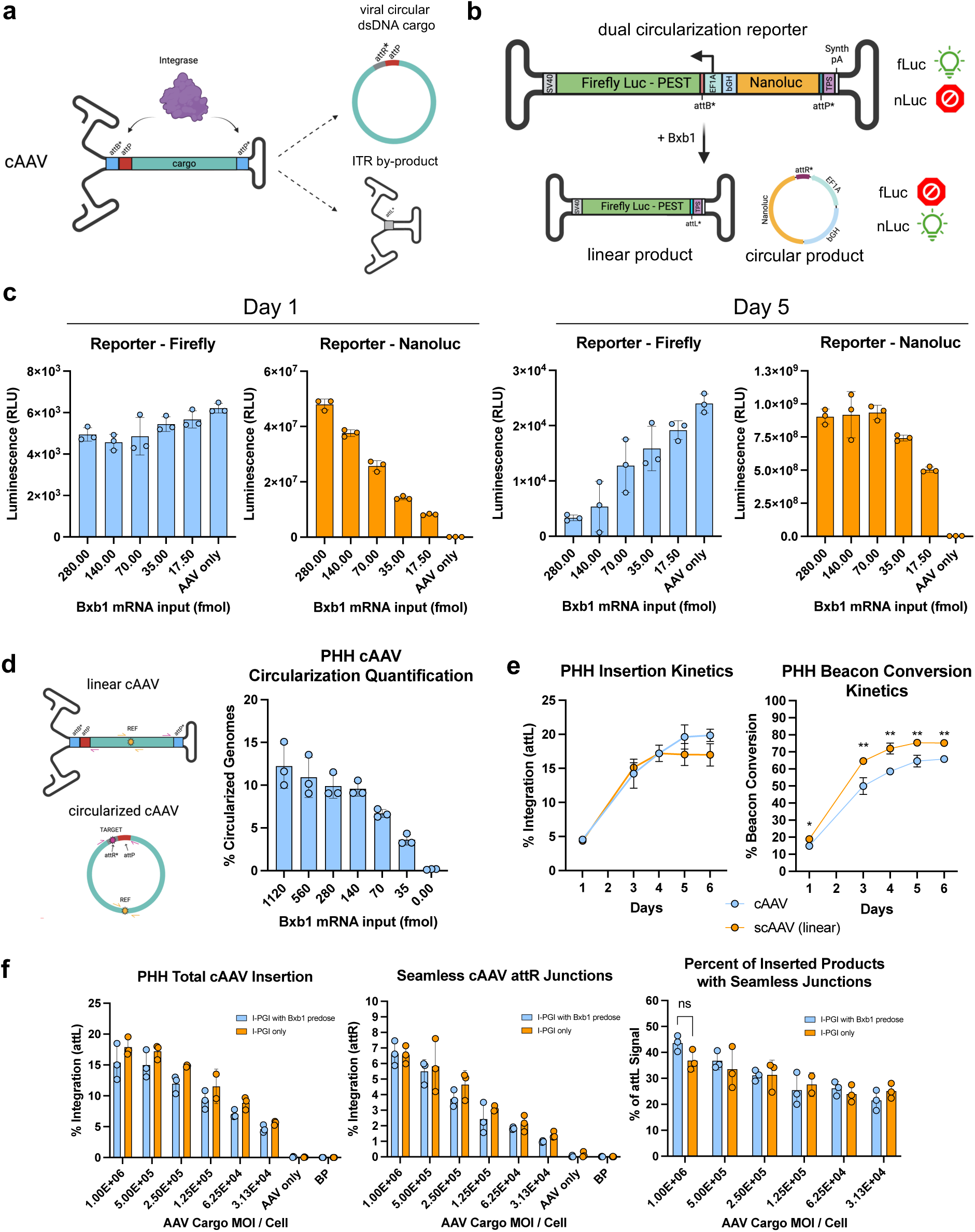
Validation of cAAV in primary human hepatocytes. (**a**) Schematic of cAAV design and mechanism. Orthogonal attachment sites (attB*, attP*) are added to an attP AAV genome to facilitate Bxb1-mediated intramolecular recombination. The reaction produces a circular dsDNA attP cargo and an ITR by-product. (**b**) Design of a ssAAV dual luciferase circularization reporter to validate cAAV function. In linear native format, EF1A promoter expresses Firefly luciferase and Nanoluc is not expressed. When circularized, EF1A promoter expresses Nanoluc in the circular episome and Firefly luciferase is not expressed in the ITR by-product. (**c**) Relative amounts of linear or circularized cAAV reporter in PHH on days 1 and 5 following Bxb1 mRNA transfection. (**d**) Absolute quantification of cAAV circularization frequency on day 5 following Bxb1 mRNA transfection in PHH. (**e**) I-PGI outcome with cAAV or scAAV cargo over a 6-day time course in PHH. Statistical significance was calculated using Student’s unpaired two tailed t-test (ns indicates not significant, * P<0.05, ** P<0.01). (**f**) The effect of staggering cargo circularization and I-PGI on seamless insertion junction outcome in PHH. Predose is Bxb1 mRNA transfected twice, on day 2 and on day 5. For **e-f**, I-PGI was performed at the *PAH* locus through a single transfection of synthetic atgRNA, nCas9-RT and Bxb1 mRNAs. Data reflects the mean and standard deviation of 3 biological replicates.

The effect of attB* and attP* orientation does not impact the efficiency of circle production in HepG2 (Extended Data Fig. 4). Because attP is present in the 5’ end of the cargo sequence for gene insertion, we used a 5’ attB* and 3’ attP* orientation to avoid highly homologous flanking sequences in the cargo. The distance between 5’ attB* and attP (for integration) was optimized to minimize interference from potential spatial constraints during integrase binding. Using purified Bxb1 and synthetic dsDNA substrates in a biochemical recombination reaction, we observed no differences in activity when moving the attB* and attP from 0 to 100 base pairs apart (Extended Data Fig. 5). All subsequent cAAV insertion experiments were performed with scAAV genome type containing adjacent attB*-attP to retain space within the cargo and avoid unnecessary sequences.

### Seamless Integration with cAAV

cAAV is a functional cargo in hepatocytes and genome circularization can occur simultaneously with I-PGI without compromising DSB-free integration efficiencies. With I-PGI, cAAV integrated into about 20% of alleles in PHH as measured by ddPCR. However, cAAV exhibited a slight reduction in cargo potency compared to linear WT scAAV, as indicated by the conversion of attB to attL (Fig. 2e). Given that integration of cAAV as linear or circularized DNA would produce identical attL junctions, we focused on attR junctions, which can distinguish between both insertion products (Extended Data Fig. 6). To determine levels of seamless insertion, we compared seamless attR junctions to total insertion by attL. We observed seamless cAAV insertion junctions in about 6.5% of alleles following I-PGI in PHH, representing ∼40% of total integration products (attL), with no significant differences from Bxb1 predosing (Fig. 2f). Cargo circularization by integrases was also used to insert longer sequences with a 30-kilobase helper-dependent adenovirus, demonstrating that this approach is effective for large linear cargo (Extended Data Fig. 7).

We then integrated cAAV into preinstalled beacons at the mouse Rosa26 locus to determine seamless insertion outcome *in vivo*. Rosa26 attB mice were treated with either linear, WT scAAV or cAAV seven days prior to LNP Bxb1 mRNA dosing, and livers were harvested for insertion analysis after an additional seven days (Fig. 3a). We observed an average total allelic integration of 35% with cAAV, marginally lower than that of WT scAAV (40%) (Fig. 3b). We examined seamless attR junctions as described previously and found an average of 32% allelic insertion of circularized cAAV, demonstrating that ∼90% of all insertion events contained DSB-free junctions (Fig. 3c). Using ONT long-read sequencing, cAAV showed homogenous seamless insertion structures with full alignment at the attR junction (Fig. 3d-e; Extended Data Fig. 8). WT scAAV exhibited DSB-dependent insertion structures, with some evidence of infrequent deletions spanning into the cargo sequence, as observed previously in PHH (Fig. 3f; Extended Data Fig. 9). While most cAAV insertion events mapped to a DSB-free outcome, we did observe a low abundance of reads mapping to linear integration structures with a typical DSB repair signature (Table 1; Extended Data Fig. 10). Although not quantitative, the high abundance of seamless integration reads supports our ddPCR findings, suggesting that cAAV is a robust cargo *in vivo* that enables significant levels of DSB-free gene insertion via Bxb1 (Fig. 3g).

**Figure 3.**
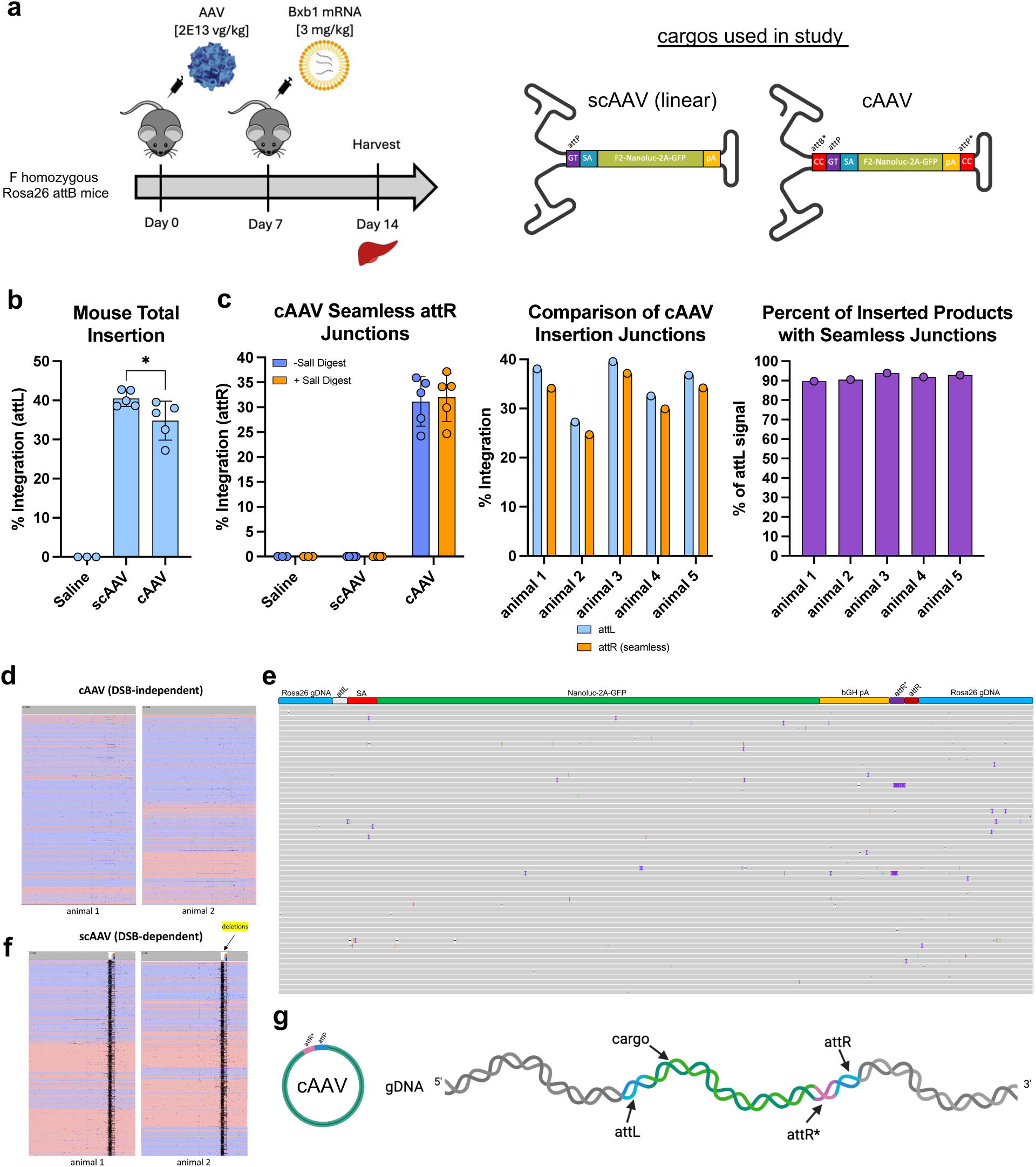
Bxb1-mediated seamless insertion with cAAV in vivo. (**a**) Schematic overview of study design for comparing insertion efficiencies and structures of scAAV and cAAV in mice. scAAV cargo contains the sequence of cAAV without orthogonal attachment sites (CC dinucleotide). (**b**) Quantification of total insertion (attL) efficiencies with scAAV and cAAV at the mouse *Rosa26* locus. Each data point represents a single animal (n=5 for scAAV and cAAV, n=3 for control). Statistical significance was calculated using Student’s unpaired two tailed t-test (* P = 0.047). (**c**) Comparison of total (attL) and seamless (attR) insertion junctions in the cAAV group. (**d**) Representative long-read sequencing alignments of seamless cAAV insertion structures from 2 animals. (**e**) Representative expanded view of cAAV seamless integration alignment from one animal where each line represents an individual read with alignment (grey). (**f**) Representative long-read sequencing alignments of scAAV insertion structures from 2 animals. (**g**) Graphic illustration of seamless insertion outcome with cAAV. For **b-c**, data reflects the mean and standard deviation unless displayed as individual animals. For **d** and **f**, each plot represents collapsed read alignment from one animal in the forward (red) or reverse (blue) strand orientation. Deletions are shown in black.

**Table 1.**
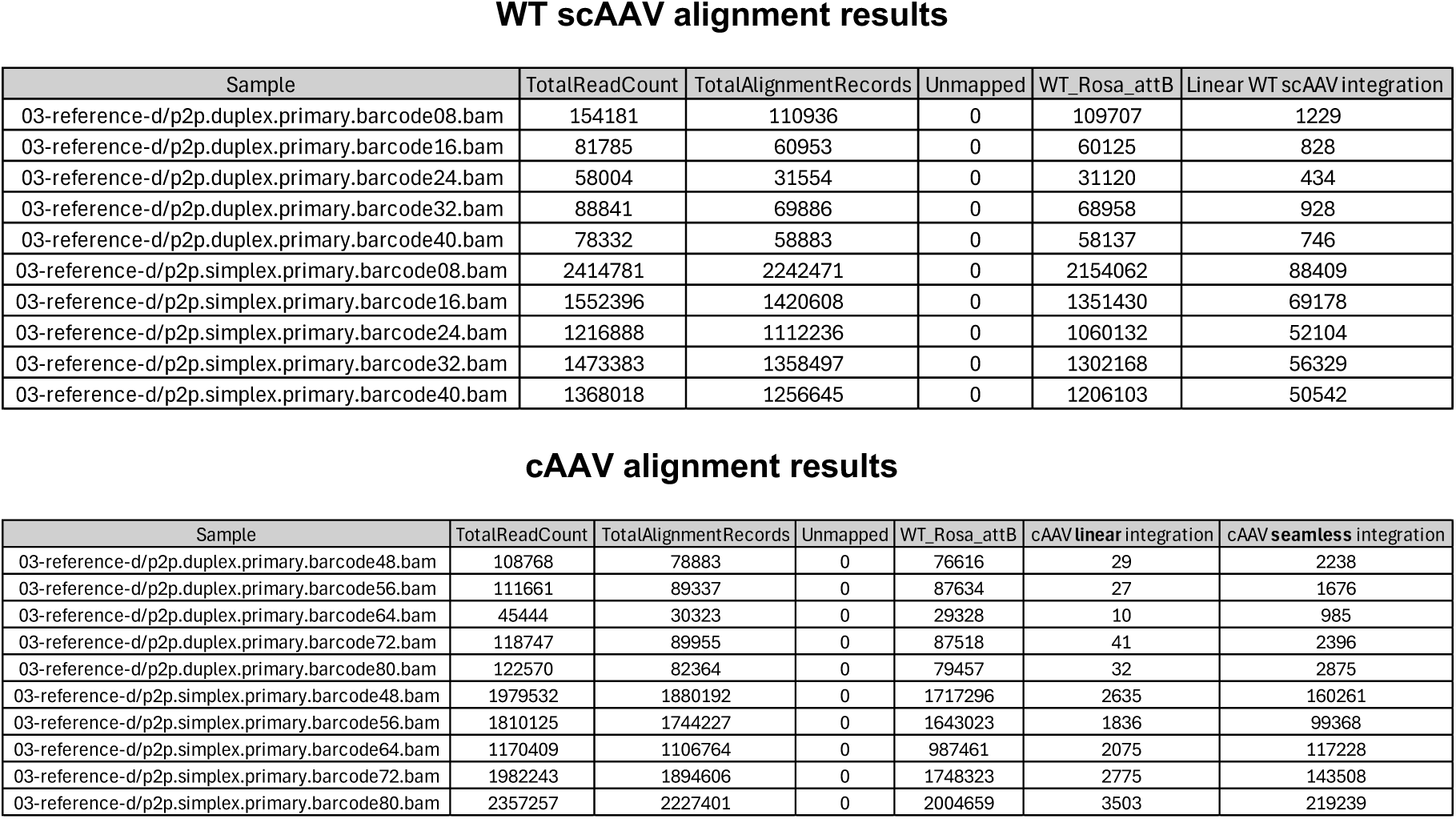
Nanopore read mapping from cAAV in vivo study. Processed reads were aligned to beacon placement or integration references for each respective cargo. Total mapped insertion read numbers are reflected under each cargo reference for each animal (barcode).

WT scAAV alignment results
| Sample | TotalReadCount | TotalAlignmentRecords | Unmapped | WT_Rosa_attB | Linear WT scAAV integration |
| --- | --- | --- | --- | --- | --- |
| 03-reference-d/p2p.duplex.primary.barcode08.bam | 154181 | 110936 | 0 | 109707 | 1229 |
| 03-reference-d/p2p.duplex.primary.barcode16.bam | 81785 | 60953 | 0 | 60125 | 828 |
| 03-reference-d/p2p.duplex.primary.barcode24.bam | 58004 | 31554 | 0 | 31120 | 434 |
| 03-reference-d/p2p.duplex.primary.barcode32.bam | 88841 | 69886 | 0 | 68958 | 928 |
| 03-reference-d/p2p.duplex.primary.barcode40.bam | 78332 | 58883 | 0 | 58137 | 746 |
| 03-reference-d/p2p.simplex.primary.barcode08.bam | 2414781 | 2242471 | 0 | 2154062 | 88409 |
| 03-reference-d/p2p.simplex.primary.barcode16.bam | 1552396 | 1420608 | 0 | 1351430 | 69178 |
| 03-reference-d/p2p.simplex.primary.barcode24.bam | 1216888 | 1112236 | 0 | 1060132 | 52104 |
| 03-reference-d/p2p.simplex.primary.barcode32.bam | 1473383 | 1358497 | 0 | 1302168 | 56329 |
| 03-reference-d/p2p.simplex.primary.barcode40.bam | 1368018 | 1256645 | 0 | 1206103 | 50542 |

| Sample | TotalReadCount | TotalAlignmentRecords | Unmapped | WT_Rosa_attB | cAAV linear integration | cAAV seamless integration |
| --- | --- | --- | --- | --- | --- | --- |
| 03-reference-d/p2p.duplex.primary.barcode48.bam | 108768 | 78883 | 0 | 76616 | 29 | 2238 |
| 03-reference-d/p2p.duplex.primary.barcode56.bam | 111661 | 89337 | 0 | 87634 | 27 | 1676 |
| 03-reference-d/p2p.duplex.primary.barcode64.bam | 45444 | 30323 | 0 | 29328 | 10 | 985 |
| 03-reference-d/p2p.duplex.primary.barcode72.bam | 118747 | 89955 | 0 | 87518 | 41 | 2396 |
| 03-reference-d/p2p.duplex.primary.barcode80.bam | 122570 | 82364 | 0 | 79457 | 32 | 2875 |
| 03-reference-d/p2p.simplex.primary.barcode48.bam | 1979532 | 1880192 | 0 | 1717296 | 2635 | 160261 |
| 03-reference-d/p2p.simplex.primary.barcode56.bam | 1810125 | 1744227 | 0 | 1643023 | 1836 | 99368 |
| 03-reference-d/p2p.simplex.primary.barcode64.bam | 1170409 | 1106764 | 0 | 987461 | 2075 | 117228 |
| 03-reference-d/p2p.simplex.primary.barcode72.bam | 1982243 | 1894606 | 0 | 1748323 | 2775 | 143508 |
| 03-reference-d/p2p.simplex.primary.barcode80.bam | 2357257 | 2227401 | 0 | 2004659 | 3503 | 219239 |

### AAV.AD Transduces Hepatocytes

We produced AAV.AD particles containing transgenes to assess the transduction potential of PHH. In our standard ssAAV backbone, we replaced the 5’ ITR with the ITR AD domain and removed the 3’ ITR, which allows for ∼2,300 kb of cargo sequence in addition to the plasmid backbone (Fig. 4a). The packaging of AAV.AD into AAV-8 and AAV-LK03 serotypes yielded usable titers, and capsids contained a relaxed circular genome, as evidenced by Exo1 and T5 digestion (Table 2, Fig. 4b). Further, we infected HEK293 and PHH with an AAV.AD promoter-driven Nanoluc reporter and observed dose-dependent transduction (Fig. 4c). Together, these results confirm that AAV.AD can transduce PHH to facilitate delivery of a circular cargo.

**Figure 4.**
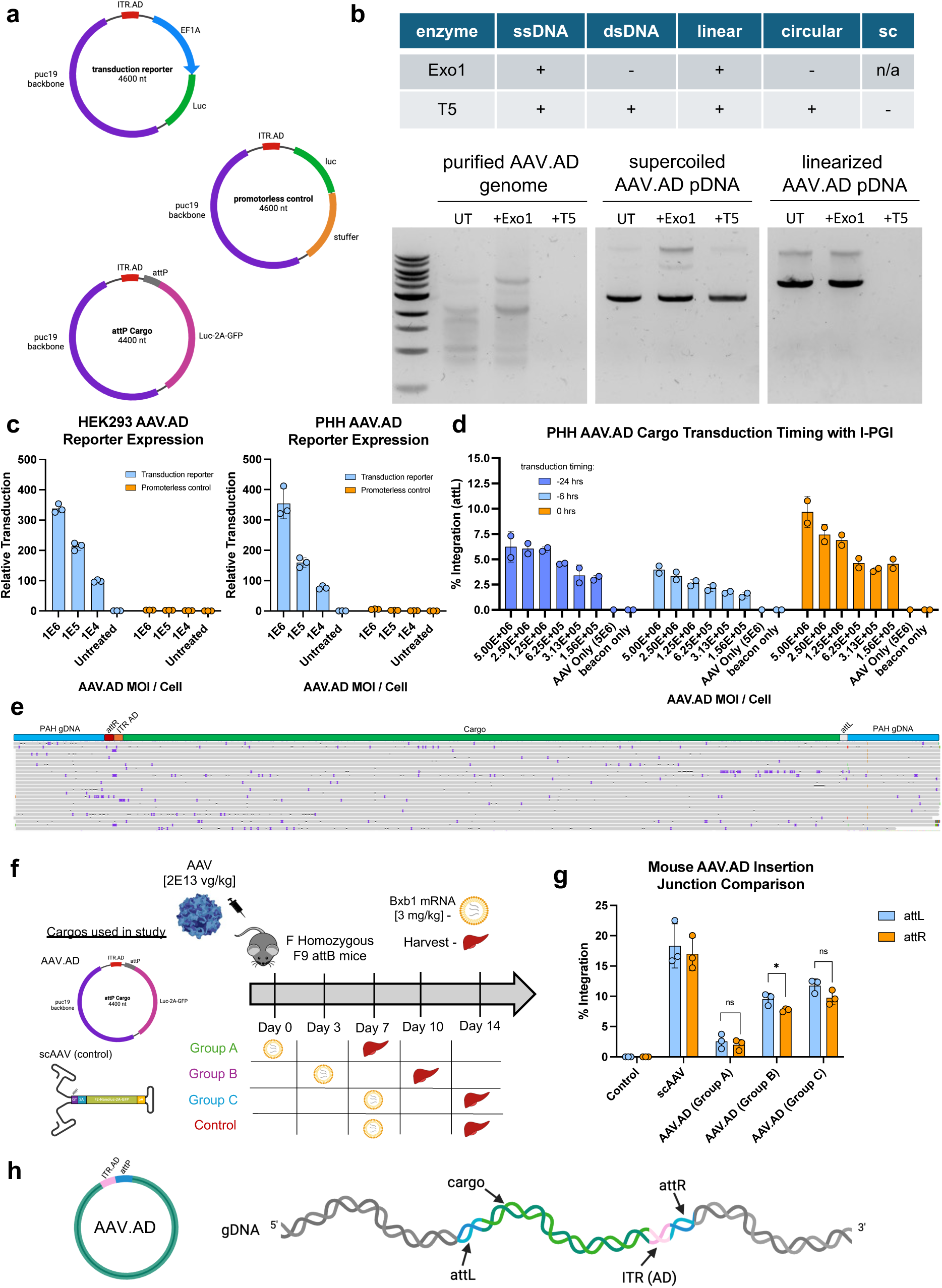
Bxb1-mediated seamless insertion with AAV.AD in PHH and mice. (**a**) Schematic of AAV.AD transfer plasmids designed for transduction reporter and integration studies. (**b**) Native agarose gel electrophoresis of AAV.AD with or without exonuclease treatment to determine DNA type. Table represents exonuclease activity on DNA substrate type including supercoiled (sc). (**c**) AAV.AD transduction efficiency in HEK293 and PHH using promoter-driven Nanoluc and promoterless control reporters. (**d**) The effect of modulating AAV.AD transduction timing on I-PGI outcome at the *PAH* locus in PHH. Transduction timing represents hours prior to single dose I-PGI transfection of synthetic atgRNAs, nCas9-RT and Bxb1 mRNAs. (**e**) Expanded view of AAV.AD seamless integration alignment in PHH where each line represents an individual read with alignment (grey). (**f**) Schematic overview of study design to evaluate insertion efficiency of AAV.AD compared to scAAV in mice. (**g**) Quantification of AAV.AD and scAAV insertion at the mouse *F9* locus (n=3 per group) where each data point reflects a single animal. Statistical significance was calculated using Student’s unpaired two tailed t-test (ns indicates not significant, * P<0.05 ). (**h**) Graphic illustration of seamless insertion outcome with AAV.AD. Data reflects the mean and standard deviation of 2-3 biological replicates. _21_

**Table 2.** Titer results from AAV.AD packaging into AAV-8 and AAV-LK03 capsids.

| Serotype | Replicate | Titer (GC/mL) |
| --- | --- | --- |
| LK03 | 1 | 1.13E+13 |
| LK03 | 2 | 3.68E+13 |
| 8 | 1 | 5.42E+12 |
| 8 | 2 | 4.39E+12 |
| 8 | 3 | 5.12E+12 |

### Seamless Integration with AAV.AD

AAV.AD particles are viable and can be utilized for targeted seamless insertion in primary hepatocytes. We adjusted the timing between AAV.AD transduction and I-PGI transfection from 24 to 48 hours, to allow for genome second-strand synthesis, and measured integration using ddPCR. Successful insertion was observed in both conditions (2-3%), with the longer predose timing demonstrating lower insertion efficiencies (Extended Data Fig. 11). By further narrowing the AAV.AD delivery window to coincide with I-PGI transfection, we achieved up to 10% allelic integration (Fig. 4d). We then resolved the integrated structure of AAV.AD in PHH. ONT long-read sequencing showed seamless structures with no indication of DSB repair, as observed previously with linear WT AAV (Fig. 4e). Like cAAV, these findings indicate that AAV.AD is a functional circular cargo for I-PGI, achieving relevant levels of genomic integration and seamless insertion outcome in primary hepatocytes.

AAV.AD efficiently integrated into preinstalled beacons at the mouse F9 locus *in vivo*. We tested AAV.AD efficacy in female homozygous F9 attB mice with varying cargo predose timing. Mice were transduced with AAV.AD on day 0, while LNP-Bxb1 mRNA was injected on day 0 (Group A), day 3 (Group B), or day 7 (Group C) (Fig. 4f). We detected AAV.AD allelic attL insertion averages of 2.5% (Group A), 9.5% (Group B), and 11.8% (Group C), demonstrating that longer incubation times improved AAV.AD integration *in vivo*. Similar efficiencies were mostly observed at the attR junction, suggesting uniform insertion from both cargo ends. However, WT scAAV integrated ∼1.5X greater than the best AAV.AD condition, indicating potency differences between cargo types (Fig. 4g). These results demonstrate that AAV.AD is a viable cargo for *in vivo* seamless insertion in the liver (Fig. 4h).

We have developed two circular AAV cargo approaches for integrase-mediated gene insertion that are functional in human hepatocytes, can facilitate DSB-independent insertion and show efficacy *in vivo* (Extended Data Fig. 12).

## Discussion

To date, strategies for targeted gene insertion in the liver utilize programmable nucleases like Cas9 or zinc finger to facilitate homology-independent insertion of AAV through host DNA end joining mechanisms [10, 41]. Integrase-mediated programmable genomic integration (I-PGI) provides a solution for liver gene insertion through targeted seamless knock-in of cargo. When applying I-PGI to the liver, we showed that recombination with linear, wildtype self-complimentary AAV (scAAV) resulted in DNA double-stranded break (DSB) repair, despite its capability to form circular episomes during transduction [36, 37]. We speculate that these integrase-mediated DSBs are likely qualitatively different than traditional DSBs generated by nucleases (e.g. Cas9, ZFN, megaTAL). When DNA is cleaved by nucleases, often the DSB is repaired perfectly, leading to multiple rounds of cutting and repair before the editing event (e.g. indel, or capture of AAV genome) stops the cycle of continual DSB generation. In the case of large serine integrase insertion of linear genomes, there is only a single DSB event, and this event only happens upon successful integration. When analyzing these integration structures, we did observe deletions spanning into the cargo polyadenylation signal, though we have not seen evidence of transcription readthrough in our RNA-seq studies (data not shown). To circumvent potential risks associated with scAAV DSB-dependent insertion outcomes [11, 12], we developed circle-AAV (cAAV) and AAV.AD cargos that result in seamless insertion as seen through long-read sequencing.

In this study, we provided evidence that cAAV is a modular cargo that can circularize with either AAV genome type and several recombinases. Our circularization approach is also functional with a 30 kilobase adenovirus format, which allows seamless insertion of large transgenes. The production of circular DNA episomes via cAAV eliminates inverted terminal repeats (ITRs), which are known to promote genome concatemerization and random insertion *in vivo* [36–38, 42]. Therefore, we predict that cAAV may offer a more uniform cargo profile for targeted gene insertion. While we were only able to quantify episomal circularization frequency of 10-15% in PHH, this was contradicted by the circularization reporter and seamless integration quantification findings. This discrepancy may stem from contamination by linear genomes due to high virus concentrations *in vitro* and possibly from non-uncoated nuclear-localized genomes [43]. We believe the actual circularization frequency of cAAV is likely much higher, as our mouse study showed 90% of integration events had seamless junctions. We observed slight reductions in beacon conversion with cAAV and circle helper-dependent adenovirus compared to linear WT genomes, likely due to Bxb1 consumption from multiple reactions as can be observed in the biochemical recombination reaction. However, this could be mitigated using two different integrases for circularization and genomic integration. Even with Bxb1 multiplexing, allelic integration with cAAV in PHH (20%) and mice (35%) exceeds curative levels for many monogenic disorders, including phenylketonuria (PKU) [44].

We cannot formally exclude the possibility that we are measuring linear cAAV genomic integration followed by ITR excision through attB* and attP*, as the recombination products would resemble seamless structures. However, our PHH results comparing sequential and simultaneous circularization showed no significant difference in seamless integration outcomes, suggesting that this is likely not a major concern. While we did not screen alternative combinations of Bxb1 dinucleotide pairs for circularization and I-PGI, it’s possible that the kinetics of either process could be modulated using a “fast” and “slow” dinucleotide combination. This approach might reduce the low amounts of linear cAAV we observed in mice (∼10%) by allowing circularization to occur at a faster rate within a single dose condition [17, 21]. In addition, we did observe reduced seamless integration outcome in PHH compared to mouse (∼40 vs ∼90% of total insertion, respectively), likely due to conditions that do not replicate *in vivo* concentrations of cargo and integrase components [45].

While we did not investigate intermolecular Bxb1-mediated recombination events with cAAV, we anticipate that Bxb1-mediated recombination would favor intramolecular reactions due to the proximal localization of attB* and attP* within the AAV genome. This concept is also supported by our PHH results where cargo circularization was efficient in the presence of genomic integration. However, if cAAV were to undergo concatemerization through intermolecular recombination and integrate with I-PGI, transcription would only occur from a single coding sequence due to the cargo polyadenylation signal.

Alternatively, AAV.AD offers reduced complexity and may be a more effective path to achieving seamless insertion in the liver. We determined that AAV.AD is a packaged circular viral genome, consistent with previous reports [33], that retains the ability to transduce the liver like WT AAV. We achieved up to 10% seamless integration in PHH by optimizing vector dose and reducing transduction timing. In contrast, longer transduction times were necessary for *in vivo* integration, yielding a maximum allelic insertion of 12%. Although we did not confirm that AAV.AD is single-stranded, the *in vivo* transduction kinetics of single-stranded AAV may explain the discrepancies observed between studies [35, 46]. Additionally, the observed potency differences between scAAV and AAV.AD support this theory, suggesting that higher AAV.AD doses and extended transduction timing might enhance insertion efficiencies *in vivo*.

This work represents a significant first step in addressing an unreported challenge for *in vivo* integrase-mediated DSB-free gene insertion. We foresee that both cAAV and AAV.AD could provide more predictable insertion outcomes than WT AAV and be effective in tissues beyond the liver, given the ability to dictate AAV tropism through serotype selection. Future studies could explore other natural circular DNA viruses, such as anellovirus and SV40, to assess their potential for achieving DSB-free recombination outcomes like AAV.AD and cAAV. Overcoming the delivery challenge of non-viral DNA to the liver would also enable use of other circular cargos, like minicircle and nanoplasmid.

## Methods

### In vitro transcription of mRNA

All mRNAs were generated via in vitro transcription (IVT) reactions using the HiScribe T7 High Yield RNA Synthesis Kit (New England Biolabs E2040S). Coding sequences were ordered as gBlocks (IDT) and cloned into an IVT vector that contains a single copy of the 5’UTR and two copies of the 3’UTR from the human beta globin gene, in addition to a 152nt polyA tail. Plasmid DNA containing coding sequences were linearized using an XbaI restriction site located immediately downstream of the polyA tail. Linearized plasmids were then purified via phenol:chloroform extraction followed by ethanol precipitation. mRNAs were produced via IVT reactions that contain Uridine-Triphosphate or N1-Methylpseudouridine-5’-Triphosphate (TriLink BioTech), and capped co-transcriptionally with CleanCap Reagent AG (3’ OMe) (TriLink BioTech). IVT reactions were incubated at 37°C for 2 hours, followed by DNAse I digestion of the template DNA. mRNA products were purified using LiCl precipitation, quantified (Qubit Fluorometric Quantification; ThermoFisher), and checked for integrity by denaturing gel electrophoresis.

### RNA oligonucleotides

atgRNAs with custom modifications were ordered from IDT and purified in house through HPLC. Purified atgRNAs were diluted in water prior to transfection.

### LNP formulation

Lipid nanoparticles were formulated using the Precision Nanosystems Ignite. Nucleic acid payloads were diluted in a 50 mM pH 4.5 acetate buffer. All lipids were purchased from commercial vendors: ALC-0315 (Broadpharm), DSPC (NOF America), cholesterol (Avanti), and DMG-PEG2000 (NOF America). Lipids were diluted in ethanol and combined to create a final stock solution according to the desired molar ratio. Mixing was performed to achieve a final LNP composition with an N/P ratio=6. LNP were then diluted 1:1 with PBS and dialyzed overnight. LNP were concentrated using ultracentrifugation, diluted to the desired concentration using Tris-sucrose buffer pH 8.0, and sterile filtered prior to freezing. All LNP were analyzed using DLS and Ribogreen to assess size, polydispersity, and encapsulation efficiency of cargo and determined to meet the desired QC parameters.

### AAV vector design

ssAAV, scAAV and AAV.AD backbones were custom synthesized and cloned at Genscript. ssAAV backbones contained WT AAV-2 ITRs. scAAV backbones contained 5’ AAV-2 ITR and a 3’ AAV-2 ITR with the delta TRS mutation. AAV.AD backbones contained one ITR AD domain. Cargos sequences were synthesized and cloned at Genscript. All attachment sites for I-PGI contained GT dinucleotides. All attachment sites for Bxb1-mediated circularization contained either CC, AC or AG dinucleotides. The WT linear scAAV used in PHH contained an attP, splice acceptor, PAH exons 2-13 coding sequence and a bGH polyadenylation signal. The dual luciferase circularization reporter was a single-stranded genome type and constructed in the reverse orientation to contain attP*, Nanoluc (non-secreted, Promega) with a bGH polyadenylation signal, an EF1A core promoter followed by attB* and Firefly luciferase-PEST (Promega) with an SV40 polyadenylation signal. A transcription pause site and synthetic polyadenylation signal flanked nanoluc in the linear format and firefly in the circularized product to prevent ITR-based transcription. All single luciferase reporters were scAAV genome type and contained flanking attB*/attP* sites, secreted Nanoluc (Promega) followed by an EF1A core promoter. The cAAV cargo used for PHH studies was a scAAV genome type and contained an attP, splice acceptor, Nanoluc-2A-GFP coding sequence followed by bGH polyadenylation signal. This cargo was also used in the mouse studies. The linear WT scAAV control in the mouse study was the exact sequence as the cAAV cargo but lacked the additional attachment sites for circularization. The Cre and FLPe cAAV cargos were ssAAV genome type and contained either flanking LoxP sites (Cre) or FRT sites (FLPe) with an attP, splice acceptor, Nanoluc-2a-GFP coding sequence, and a bGH polyadenylation signal. The AAV.AD transduction reporter contained an EF1A promoter followed by secreted Nanoluc. The control lacked a promoter and utilized a stuffer to maintain equal genome lengths. The AAV.AD cargo used in PHH and mice contained an attP, splice acceptor, Nanoluc-2a-gfp coding sequence and a bGH polyadenylation signal. All AAV.AD genomes had a maximal size of 4,600 nucleotides to account for ssAAV packaging limit.

### AAV production

Recombinant AAV was produced using 293AAV (Cell Biolabs AAV-100) triple transfection with transfer plasmid, rep/cap and helper (Aldeveron SF058826) plasmids. AAV-8 (Genemed P-RC09) was produced for in vivo studies. AAV-DJ (Cell biolabs VPK-420-DJ) and AAV-LK03 (internally optimized) were produced for in vitro studies. Recombinant virus was treated with PEG and purified through Iodixanol gradient centrifugation. Resulting titers were determined using ddPCR with an ITR-based assay.

### hdAD vector design

Adenovirus serotype 5 gutless vectors were utilized for cargo production. Vector construction was performed in house. All attachment sites for I-PGI contained a GT dinucleotide. Reporter cargos contained attP, Nanoluc-2a-gfp coding sequence and a bGH polyadenylation signal. Factor 9 cargos contained an attP and the full genomic sequence of human factor 9 from intron 1 to exon 8 including the mouse Factor 9 3’ UTR. The circle hdAD cargos contained additional Bxb1 attachment sites with the AG dinucleotide. The WT linear versions did not include orthogonal bxb1 attachment sites.

### hdAD production

To produce recombinant hdAD, vectors were linearized and transfected in HEK293 116 Cre+ cells (Baylor University) followed by helper virus transduction. hdAD was propagated by serial coinfection and purified by CsCl ultracentrifugation. Resulting titer was determined by ddPCR using a genome specific assay.

### Cell culture

HepG2 cells were purchased from ATCC and cultured DMEM (Gibco 11965092) with 10% FBS (Gibco A3160501) in pre-coated Poly-D-Lysine (A3890401, Gibco) T75 flasks (Thermo Scientific 156472). Cells were dissociated with 0.25% Trypsin-EDTA (15400054, Gibco) and seeded in Collagen I coated 96 well plates (Corning 354407) at a density of 10k cells per well. HEK293 were purchased from ATCC (CRL-1573) and cultured in DMEM (Gibco 11965092) with 10% FBS (Gibco A3160501) in T75 flasks (Thermo Scientific 156472). Cells were dissociated with TrypLE Express (Gibco 12604013) and seeded in Collagen I coated 96 well plates (Corning 354407) at a density of 25k cells per well. Cryopreserved primary human hepatocytes (Thermofisher) were recovered in Cryopreserved Hepatocyte Recovery Media (Gibco CM7000) and plated at 37-42k cells per well. 6 hours post recovery, cells were washed and cultured in maintenance media (Gibco A1217601, CM400, A2737501). All cultures were maintained at 37C and 5% CO_2_.

### HepG2 study

All transfections were performed using 0.3 uL Lipofectamine MessengerMax (Invitrogen LMRNA015) and Opti-MEM (Gibco 31985062) 16-24 hours after plating. AAV-LK03 cAAV reporters were transduced at 5E5 MOI / cell simultaneously with Bxb1 mRNA transfection. Relative circularization was measured 3 days after transfection.

### HEK293 studies

All transfections and transductions were performed 16-24 hours after plating. For AAV.AD transduction, cells were infected at indicated MOIs and relative transduction efficiency was measured 3 days later. For circularization with alternative recombinases and plasmid substrate, cells were cotransfected with recombinase plasmid DNA (25-50 ng) and AAV transfer plasmid (50 ng) with lipofectamine 3000 Transfection Reagent (Invitrogen L300015) at a final volume of 0.2 uL per well. For circularization with alternative recombinases and AAV substrate, cells were transduced with AAV-DJ cAAV at 3E5 MOI / cell simultaneously with recombinase plasmid DNA transfection as mentioned above. Cells were lysed on day 3 and processed for ddPCR analysis to determine circularization frequency.

### Primary human hepatocyte studies

All transfections were performed using 0.3 uL Lipofectamine MessengerMax (Invitrogen LMRNA015) and Opti-MEM (Gibco 31985062). For all standard I-PGI experiments with AAV cargo, cells were treated with AAV-LK03 at 1E6 MOI / cell one day after plating and transfected with single lipoplex containing nCas9-RT mRNA (280 fmol), Bxb1 mRNA (190 fmol) and 2 pmol of each purified atgRNA two days after plating. Media was supplemented with dTTP/dGTP (NEB N0446S) at a final concentration of 200 uM (equimolar) in the well. For I-PGI experiments with hdAD cargo, cells were transfected for beacon placement on day 2 with 2 pmol of each purified atgRNA and 280 fmol of nCas9-RT mRNA with dNTP supplementation as described above. hdAD was transduced at 1E5 MOI / cell (unless indicated otherwise) simultaneously with Bxb1 mRNA (190 fmol) transfection. For measuring absolute circularization frequency with cAAV, cells were transduced with AAV-LK03 one day after plating and Bxb1 mRNA mRNA was transfected on day 2. For AAV.AD transduction reporters, AAV-DJ was transduced 1 day after plating at indicated amounts. For luciferase circularization reporter experiments, AAV-LK03 was transduced one day after plating and integrase mRNA was transfected at indicated amounts on day 2. Alternatively, adenovirus Bxb1 was used at 125 MOI / cell and cotransduced with AAV reporter on day 1 after plating. All experiments were taken down on day 5 and processed for ddPCR analysis or reporter expression. Any deviations in AAV cargo MOI, Bxb1 mRNA amount, Bxb1 predosing or take down time point are indicated in respective figures.

### *In vivo* mouse studies

All animal study procedures were approved by Explora BioLabs under IACUC protocol EB17-004-302. Transgenic C57BL/6J mice with a knock-in of the attB site in intron 1 of F9 or Rosa26 were generated by Biocytogen (Beijing, China) using CRISPR/Cas9. Mice were transferred to Biomere (Worcester, MA, USA) for breeding. Control linear AAV8 scAAV and/or experimental AAV-8 cAAV or AAV-8 AAV.AD cargos were intravenously injected into adult mice (∼20-46 weeks) at a dose of 2E13 at day 0. At days 0 -7, LNPs formulated with WT Bxb1 stabilized-2 or WT Bxb1 stabilized-4 mRNAs were intravenously injected into the mice at a dose of 3 mg/kg. Seven days after LNP injection, animals were euthanized, and liver tissue was collected from the median lobe from each animal and homogenized on Precellys Evolution (cat K002198-PEVO0-A.0 Combo, Bertin technologies, WA, USA).

### Reporter expression

Reporter expression was processed using the Nano-Glo Dual-Luciferase Reporter Assay System (Promega N1610), Nano-Glo Luciferase Assay System (Promega N1150) or CellTiter-Glo 2.0 Viability Assay (Promega G9242). Luminescence was read on the GloMax Discover Microplate Reader (Promega GM3000). Relative circularization or expression was determined by normalizing Nano-Glo to CellTiter-Glo.

### Cell lysis and gDNA purification

HEK293, HepG2 and PHH were lysed at indicated time points by removing cell culture medium, adding 50 uL of QuickExtract lysis buffer (LGC Biosearch Technologies QE0905T), shaking plates for 10 minutes at 700 RPM, transferring lysate to 96 well plate followed by a thermal cycle (75C 10 minutes, 98C 5 min and 4C hold) and a freeze thaw at -80C. Magnetic bead clean up with Ampure XP beads was performed to purify gDNA from resulting lysate (Beckman Coulter A63882). For absolute quantification of cAAV circularization frequency in PHH, cells were first washed 3X in PBS -/- (Gibco 70011-044) followed by cytoplasmic extraction (Nano-Glo HiBiT lytic buffer, Promega N246) and removal, an additional 3X washes in PBS and nuclei lysis with Quick Extract Lysis Buffer. Nuclear DNA was purified from nuclear lysate as mentioned above. Liver gDNA was extracted with quick-DNA/RNA MagBead kit (Zymo research R2131).

### Droplet digital polymerase chain reaction (ddPCR)

ddPCR was used to determine insertion, attB beacon placement and cAAV circularization frequencies. For insertion quantification, custom assays were developed to amplify either the left (attL) or right (attR) recombination junction using genome/cargo-specific primers and a probe that targeted the recombination scar. For beacon placement quantification, custom assays were developed to amplify over the targeted genomic region using two genome-specific primers and a probe that targeted attB. Signal from attB, attL or attR assays were normalized to custom reference assays targeting unedited regions of the target gene.

To calculate beacon conversion (percentage of attBs converted to attL), total insertion (attL) was normalized to total editing (attB+attL). Relative percentages of seamless insertion were determined by seamless attR signal normalized to total insertion (attL). To measure absolute episomal circularization frequency of cAAV, a custom reference assay targeted a region of the transgene coding sequence that would be present in both linear or circular forms and a target assay spanned the recombination junction with the probe binding to attR* or cargo sequence. For the seamless attR junction assay with cAAV, ddPCR reactions were incubated at room temp for 1 hour with 1E5 units of SalI (NEB R3138L) to facilitate digestion of linear cAAV insertion products prior to droplet generation. Probes were dual labeled with 3′-3IABkFQ and either 5′-carboxyfluorescein (FAM) for edit targets or 5′-hexachloro-fluorescein phosphoramidite (HEX) for reference. Assays were validated using gBlocks representing edit outcomes to test for both specificity and linearity. All primers, probes, and gBlocks were synthesized by IDT. Each reaction contained 12 µL of 2x ddPCR Supermix for probes (No dUTP) (Bio-Rad 1863025), 1.2 µL of each primer and probe mix to final concentration of 0.5 uM for each primer and 0.25 uM for each probe, 0.12 µL each of HindIII and Eco91I (FD0505 and FD0394, Thermo Fisher Scientific), 10-30 ng of DNA and water to a final volume of 24 µL. Droplets were generated on the AutoDG Instrument for automated droplet generation (186410, Bio-Rad). PCR amplification was performed with the following cycling parameters: initial denaturation at 95 °C for 10 min, followed by 40 cycles of denaturation at 94 °C for 30 s and combined annealing/extension step at 58-60 °C for 1 min, and a final step at 98 °C for 10 min. Data acquisition and analysis were performed on the QX200 Droplet Reader. All primers and probes used for ddPCR are listed below.

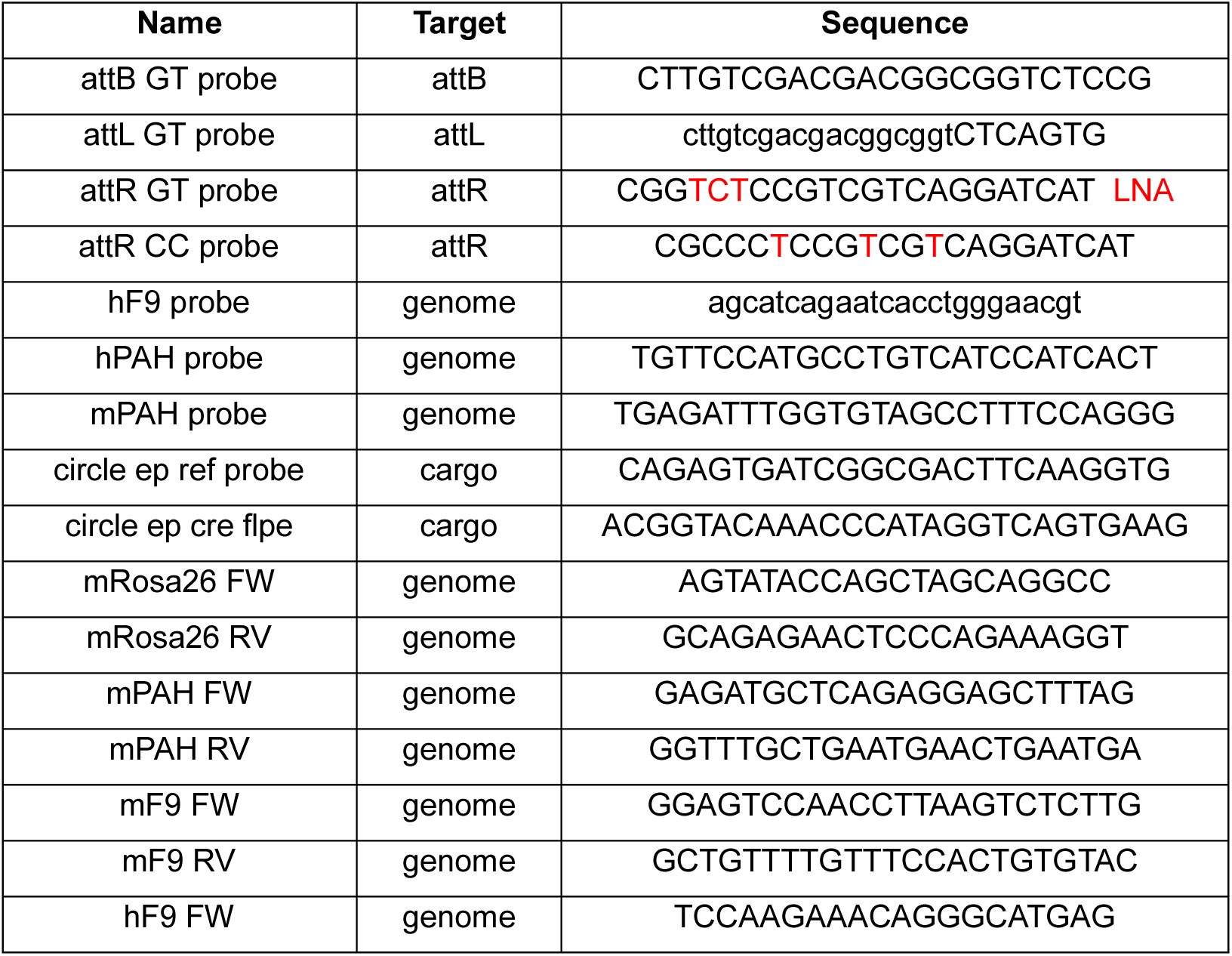
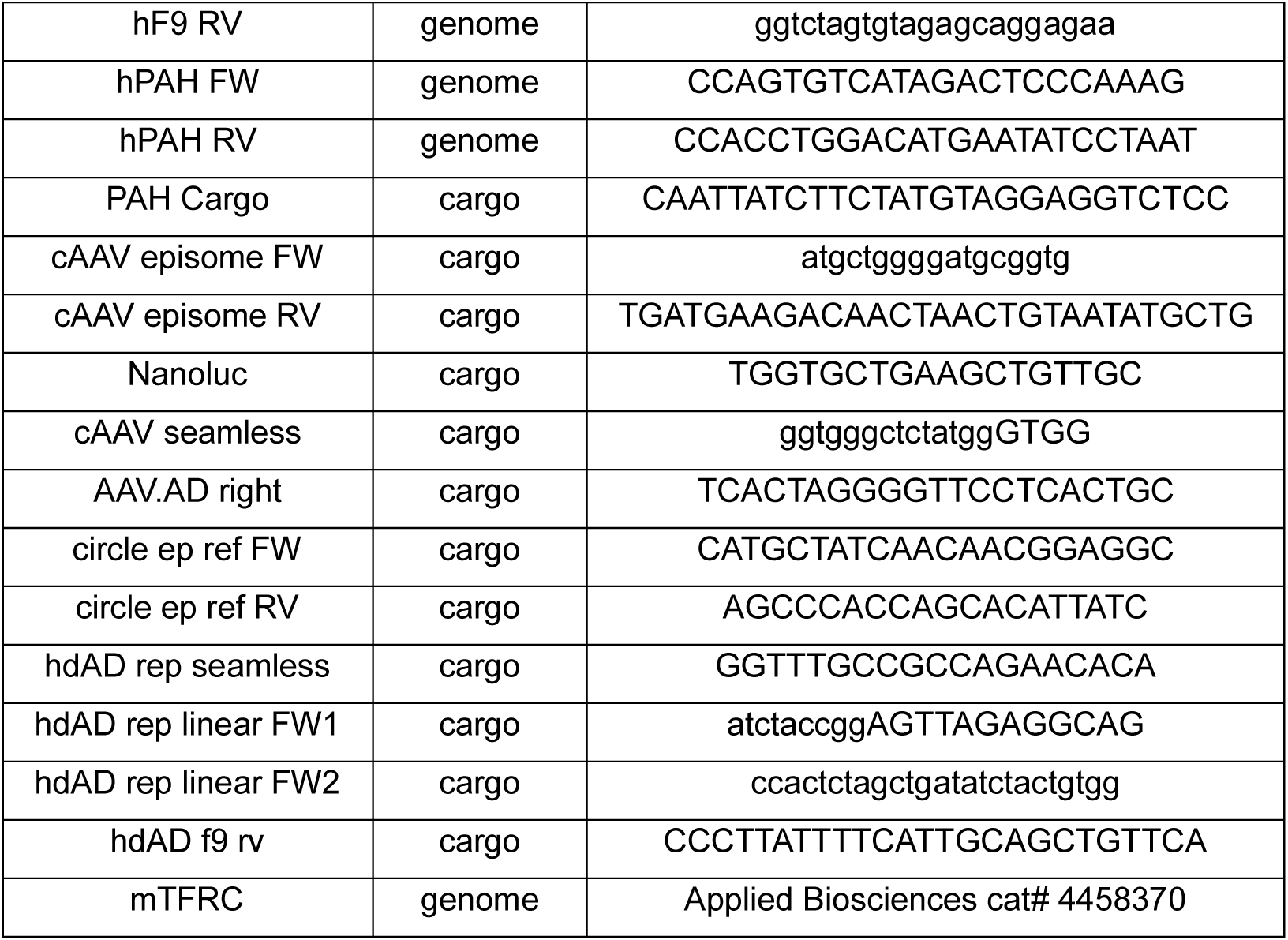

### ONT sequencing of amplified insertion products

Genomic DNA from PHH and mouse liver was prepared as described previously. The concentration of isolated genomic DNA was quantified by high sensitivity Qubit dsDNA quantification assay kit from ThermoFisher Scientific (Cat. No. Q33230). 200ng of isolated genomic DNA was used as a template for PCR amplification to amplify the targeted region of using primer sets indicated below. PrimeSTAR GXL DNA polymerase from Takara (Cat.No. R050A) was used for carrying out the PCR reaction. PCR amplification was performed with the following cycling parameters: initial denaturation at 98 °C for 3min, followed by 30 cycles of denaturation at 98 °C for 10s, annealing at 60 °C for 15s and extension step at 68 °C for 3min, and a final extension step at 68 °C for 5min. Purification of PCR amplicons are carried out using AMPure XP bead-based reagent (Beckman Coulter Cat. No. A63882) according to manufacturer’s protocol. The purified PCR amplicons were used to generate Oxford Nanopore Technology (ONT) libraries for performing long read sequencing using ONT Native Barcoding Kit 23 V14 (Cat. No. SQK-NBD114.24). The ONT libraries were generated according to manufacturer’s protocol. In brief, end prep reaction was carried out for each samples using 200fmol of purified PCR amplicons followed by native barcode ligation. For the last step of library preparation, ONT adapter was ligated to the pooled barcoded samples. 10fmol of the final pooled ONT library was loaded onto MinION Flow Cell R10.4.1 from ONT (Cat. No. FLO-MIN114) and the sequencing run was carried out for 2 days. The long read sequencing data obtained from ONT was analyzed using the in-house bioinformatics pipeline. Primers used for insertion product amplification are listed below.

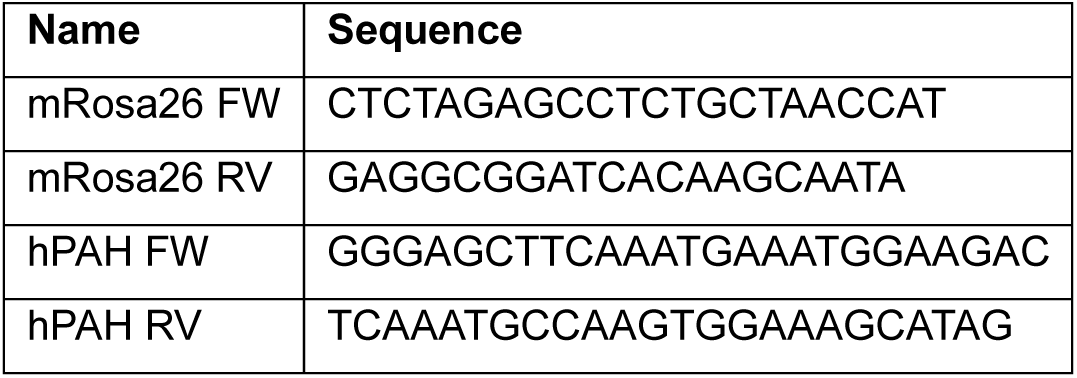

### ONT sequencing of native gDNA insertion products

The Ligation sequencing gDNA – Cas9 enrichment kit (ONT SQK-CS9109) was utilized for PCR-free sequencing of WT linear scAAV insertion structures. sgRNAs were designed to target the insertion product region using a tiling approach (2 cuts on both sides). 5 ug of genomic DNA was dephosphorylated using Quick CIP (NEB M0525S) for 30 minutes at 37C. Cas9 RNPs were formed with Alt-R S. pyogenes HiFi Cas9 nuclease V3 (IDT 1081060) and pooled sgRNAs followed by a biochemical Cas9 cleavage reaction with genomic DNA substrate to expose 5’ phosphates on the region of interest. The subsequent steps for library preparation followed the manufacture’s protocol. The final ONT library was loaded onto MinION Flow Cell R10.4.1 from ONT (Cat. No. FLO-MIN114) and the sequencing run was carried out for 2 days.

### ONT data analysis

Raw signals from nanopore were basecalled and demultiplexed using Dorado v 0.7 with the *sup* model. Adapters were trimmed using Porechop and reads were filtered with chopper to remove reads shorter than 300 bp and average quality less than 10. Read quality plots were generated by Nanoplot. Filtered reads were mapped to prebuilt reference sequence using bwa mem (-x ont2d). Data was visualized on Integrative Genomics Viewer.

https://github.com/nanoporetech/dorado

https://github.com/rrwick/Porechop

https://github.com/wdecoster/chopper

https://github.com/wdecoster/NanoPlot

### AAV.AD genome purification and digestion

1E12 AAV.AD particles were treated with 25 units of DNase1 (Thermo EN0521) for 30 min at 37C to digest any plasmid DNA within the AAV sample. Capsids were then lysed at 55C for 4 hours. Genomes were purified using 1X phenol chloroform extraction (Invitrogen 15593031), 2X chloroform wash (Sigma 366927), ethanol (Fisher scientific BP2818100) precipitation followed by DNA resuspension in water. Purification was performed in MaXtract High Density tubes (Qiagen 129056). AAV.AD genomes were then digested by Exonuclease I (NEB M0293S) or T5 Exonuclease (NEB M0663S) and run on a native agarose gel (Invitrogen 16500500) alongside untreated genomes. For additional controls, the AAV.AD transfer plasmid DNA in supercoiled (untreated) or linearized (XhoI, NEB R0146S) states were subjected to digestion by both exonucleases mentioned above and run alongside AAV.AD purified genome samples.

### Biochemical measurement of Bxb1 activity

Substrates for biochemical assays were generated using synthetic gBlocks (IDT) designed for each specific target. The size of each input and product were: attP* 201 bp, attB* 259 bp (constant), attL 127 bp and attR 459 bp. Activity assays were carried out with 20 nM of each substrate in a buffer consisting of 25 mM Tris pH 8.0, 100 mM KCl, 50 mM NaCl, 1 mM Spermidine, 5 mM MgCl*2*, 2.5 mM DTT, 5% Glycerol. Enzyme and substrate mixtures were incubated for 1 hour at 30 °C and reactions were stopped by addition of Proteinase K and incubation at 37 °C for 30 minutes. Samples were visualized using a 4200 Tapestation (Agilent, G2991BA) using D1000 DNA Screen Tapes (Agilent, 5067-5582). Area under the curve measurements for the resulting bands were used to calculate the fraction of each substrate/product.

### Other

Figure schematics were generated using Biorender and Microsoft PowerPoint. Graphs and data analysis were performed with GraphPad Prism 10. All results are reported as mean and standard deviation of biological duplicates or triplicates for *in vitro* experiments and individual results from 3-5 replicates with standard deviation for *in vivo* experiments.

---

**Extended Data Figure 1.**
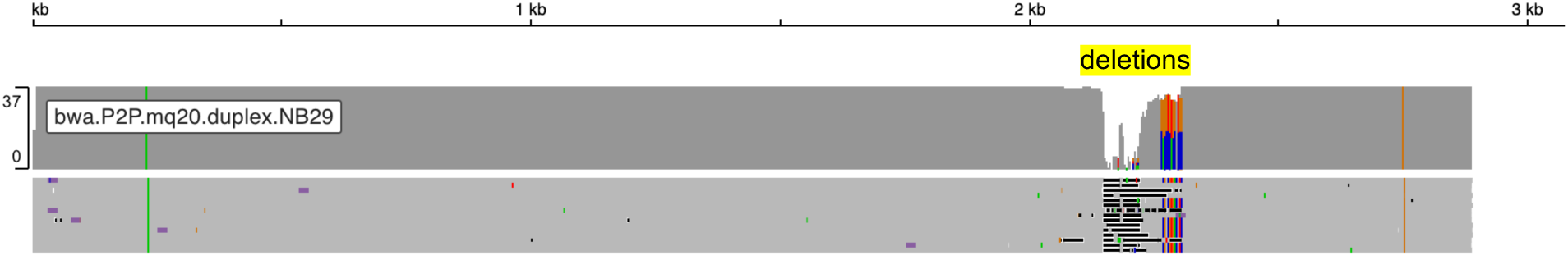
Alignment of WT scAAV insertion structures using ONT targeted native gDNA sequencing. Reads were aligned to a full-length scAAV linear insertion reference sequence. Each line represents a single read with alignment (grey) and deletions (black).

**Extended Data Figure 2.**
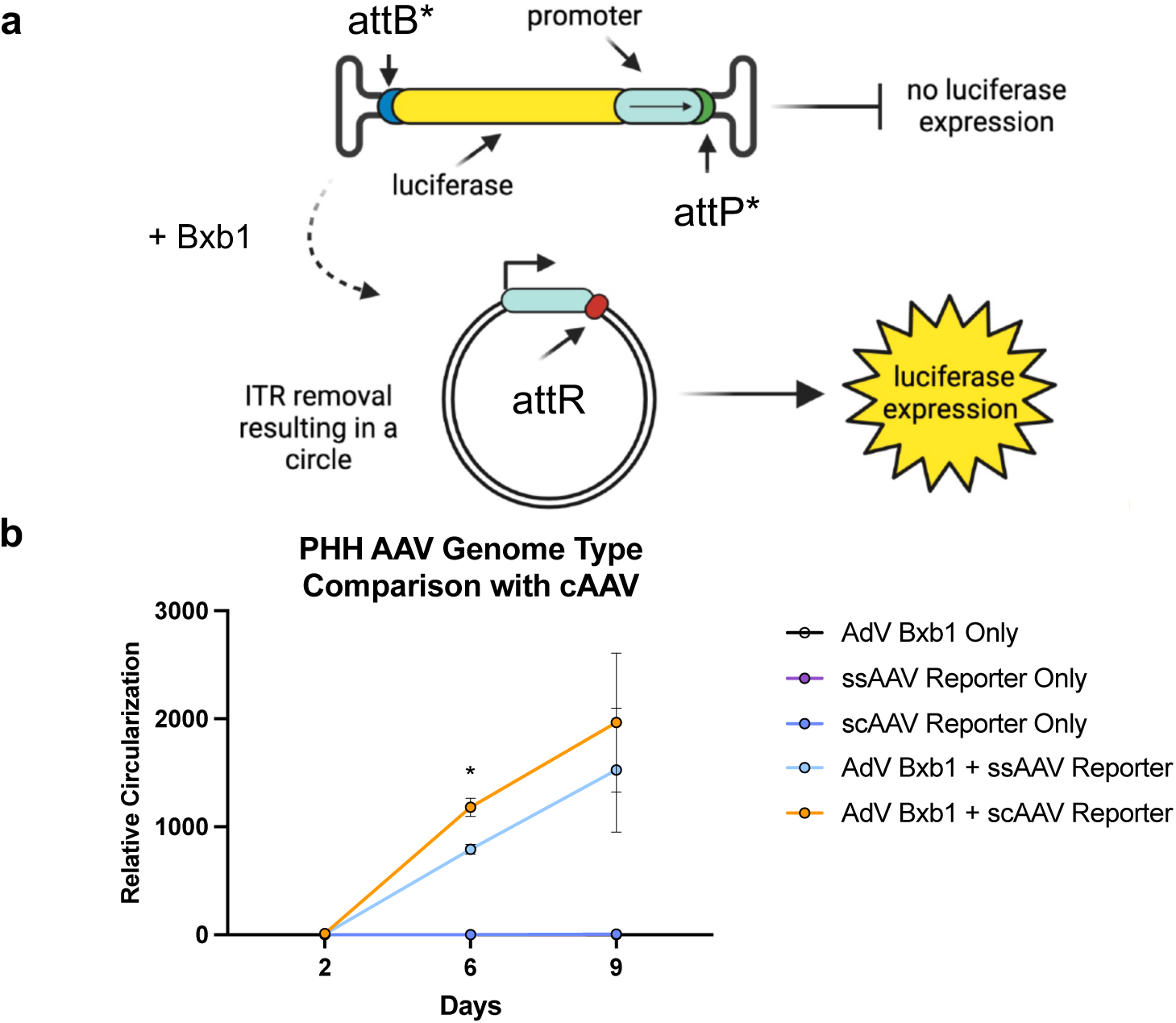
cAAV circularization kinetics in scAAV and ssAAV genome formats. (**a**) Graphic depiction of circularization reporters where luciferase is only expressed upon successful genome circularization. (**b**) Relative circularization kinetics of scAAV and ssAAV reporters with adenovirus (AdV) Bxb1 in PHH. Statistical significance was calculated using Student’s unpaired two tailed t-test (* P<0.05). Data reflects the mean and standard deviation of 2 biological replicates.

**Extended Data Figure 3.**
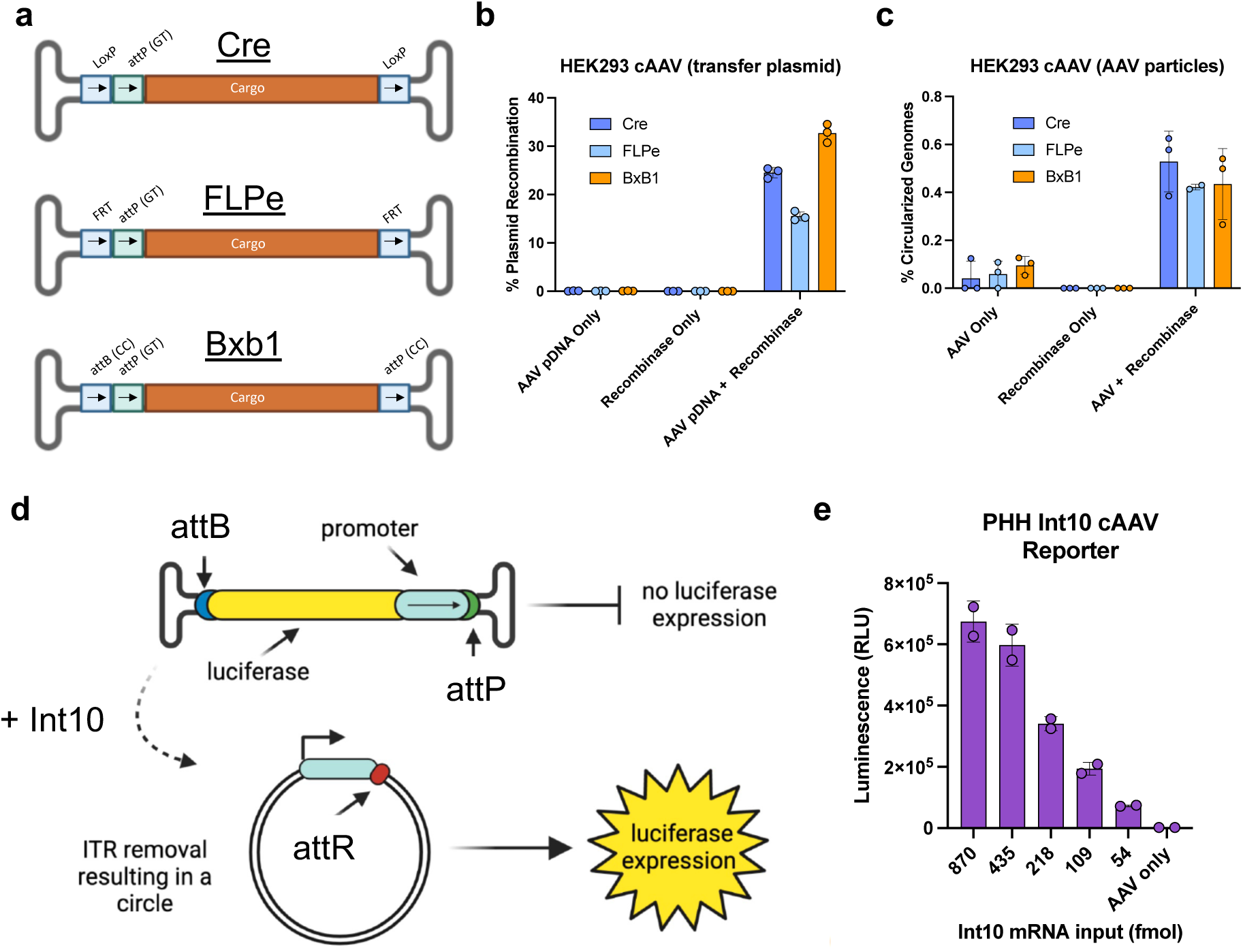
cAAV with alternative recombinases. (**a**) Graphic depiction of single-stranded cAAV genome design with either LoxP (Cre recombinase), FRT (FLPe recombinase) or attB/attP (Bxb1 integrase). Arrows indicate orientation of each recombination site. (**b**) Intramolecular recombination efficiencies with AAV plasmid DNA substrate in HEK293 using Cre, FLPe and Bxb1 recombinases. (**c**) Circularization quantification of ssAAV substrate in HEK293 using Cre, FLPe and Bxb1 recombinases. (**d**) Graphic depiction of Int10 cAAV reporter where luciferase is expressed upon successful AAV genome circularization. (**e**) Effect of Int10 mRNA dose response curve on relative cAAV circularization in PHH. Data reflects the mean and standard deviation of 2-3 biological replicates.

**Extended Data Figure 4.**
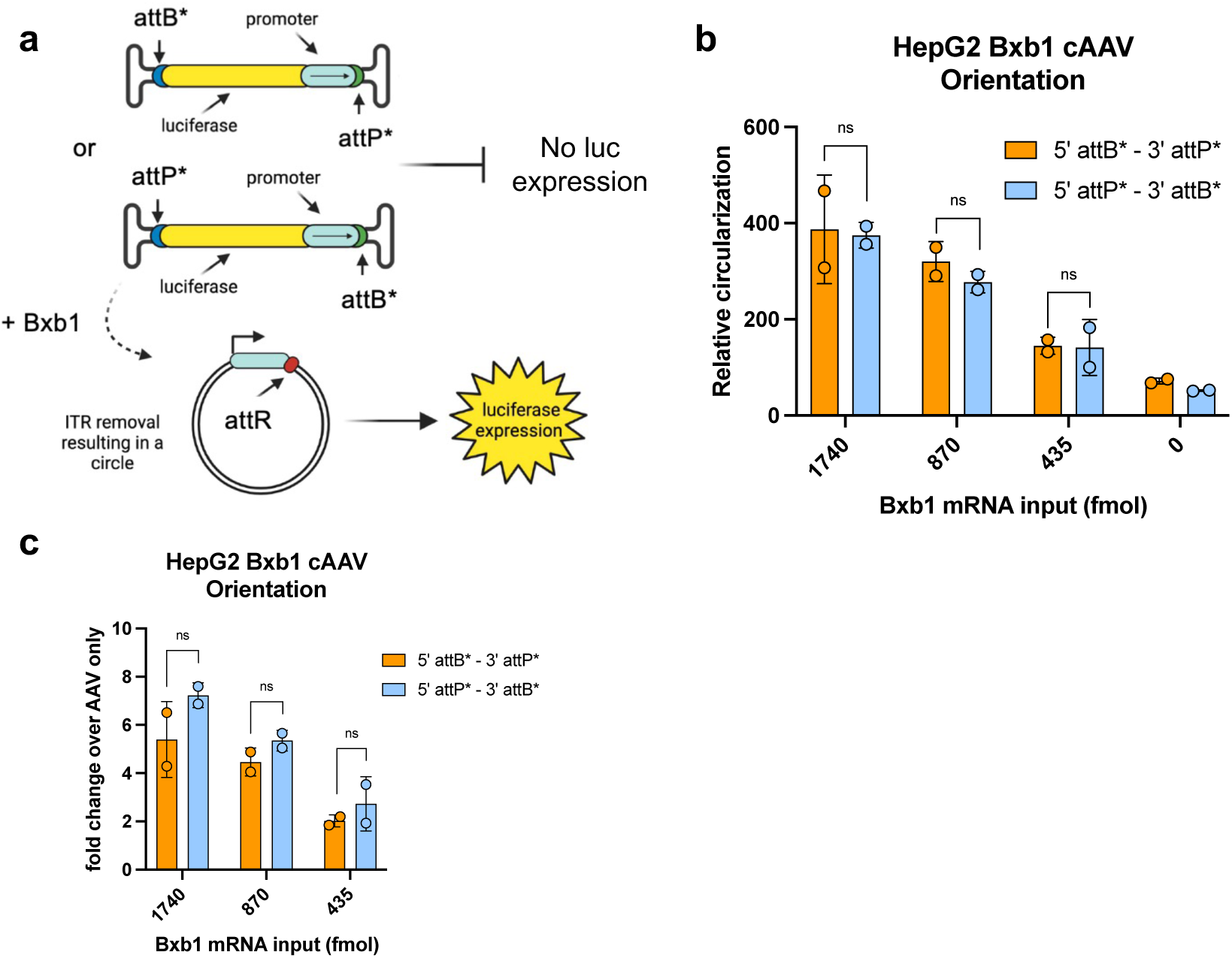
Evaluation of attB* and attP* orientation on cAAV circularization. **(a**) Graphic depiction of Bxb1 cAAV reporter design where luciferase is expressed only upon successful circularization. attB* and attP* are placed at the 5’ and 3’ ends in both combinations. (**b**) Comparison of relative circularization efficiencies between both attachment site orientation combinations in HepG2. (**c**) Fold-change in relative circularization over AAV only control from **b**. Statistical significance was calculated using Student’s unpaired two tailed t-test (ns indicates not significant). Data reflects the mean and standard deviation of 2 biological replicates.

**Extended Data Figure 5.**
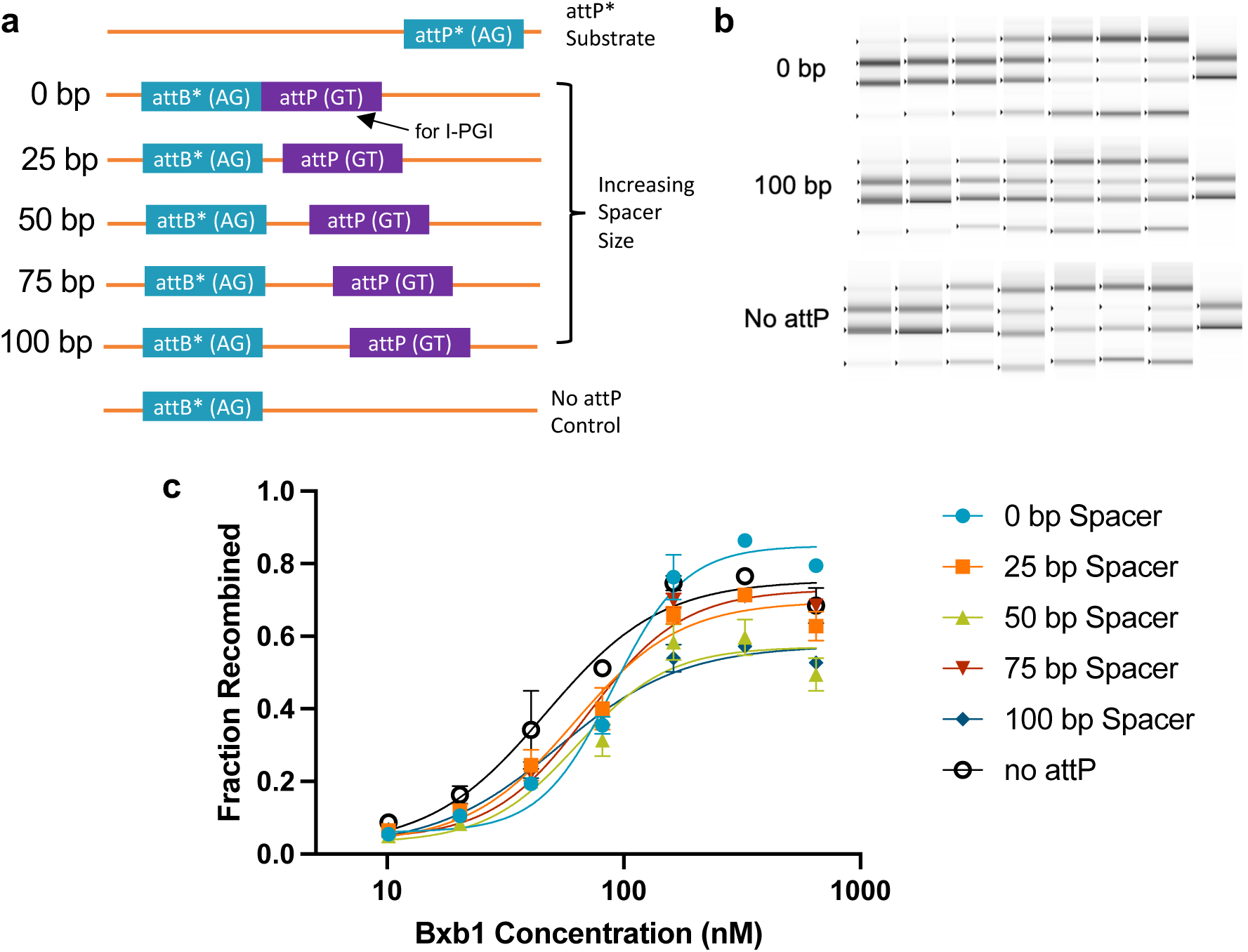
Biochemical optimization of cAAV attB* - attP distance. (**a**) Schematic representation of Bxb1 substrates designed for biochemical recombination. (**b**) Representative tapestation images of resulting recombination products. (**c**) Biochemical recombination efficiency for each attB* substrate with attP* substrate.

**Extended Data Figure 6.**
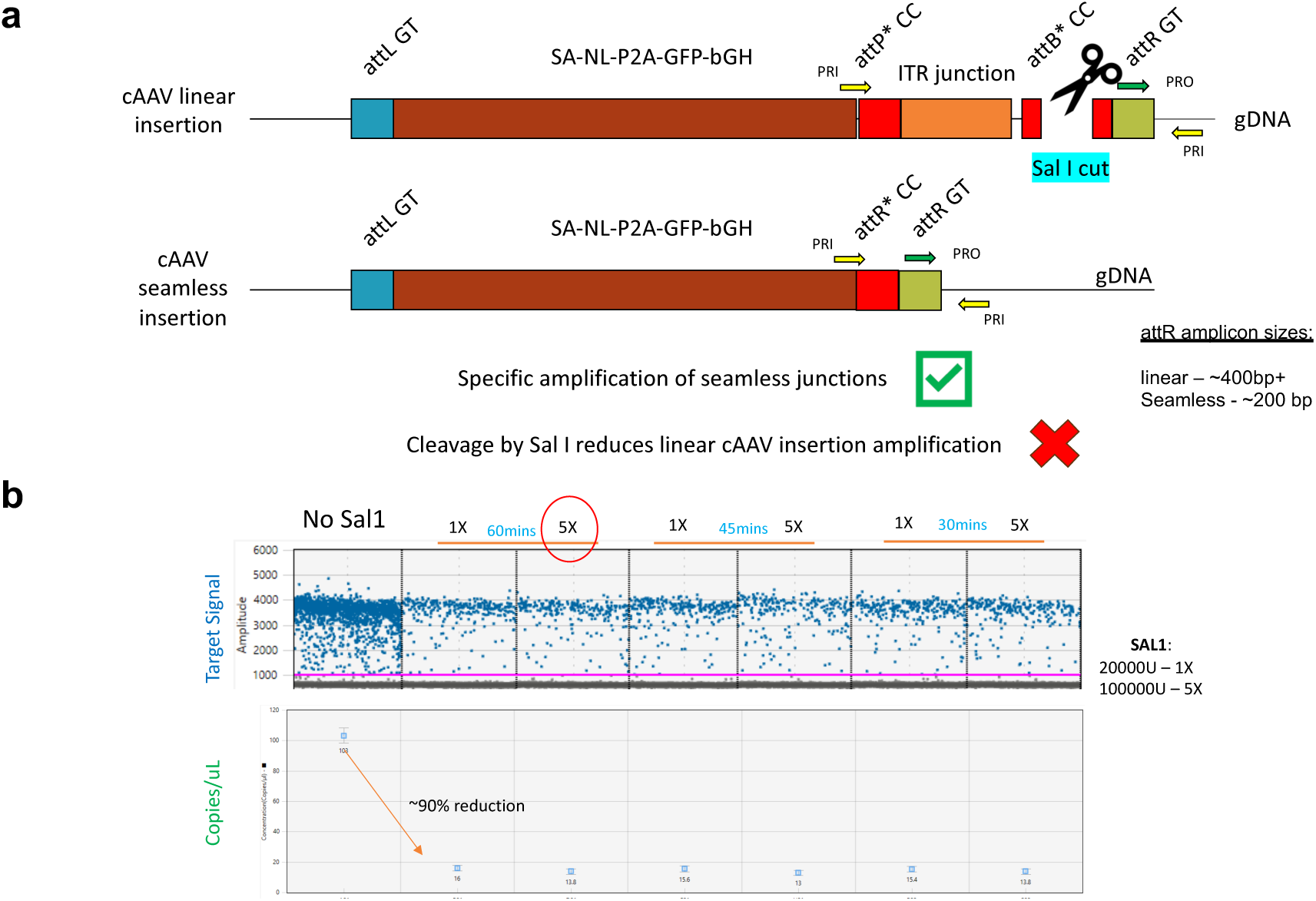
Design and validation of a cAAV seamless attR ddPCR assay. (**a**) Schematic overview of assay concept. cAAV can integrate in linear format (non-circularized) or circular format (seamless). Primers (PRI) and probes (PRO) bind to both structures (100% homology) which can lead to amplification of linear products. Digestion of DNA with Sal I restriction enzyme specifically cleaves attB* in linear cAAV insertion structures to prevent amplification. (**b**) Validation of approach and optimization of SalI digest conditions. 100,000 units of SalI and 60-minute digestion length were chosen to ensure saturating digestion conditions with gDNA samples.

**Extended Data Figure 7.**
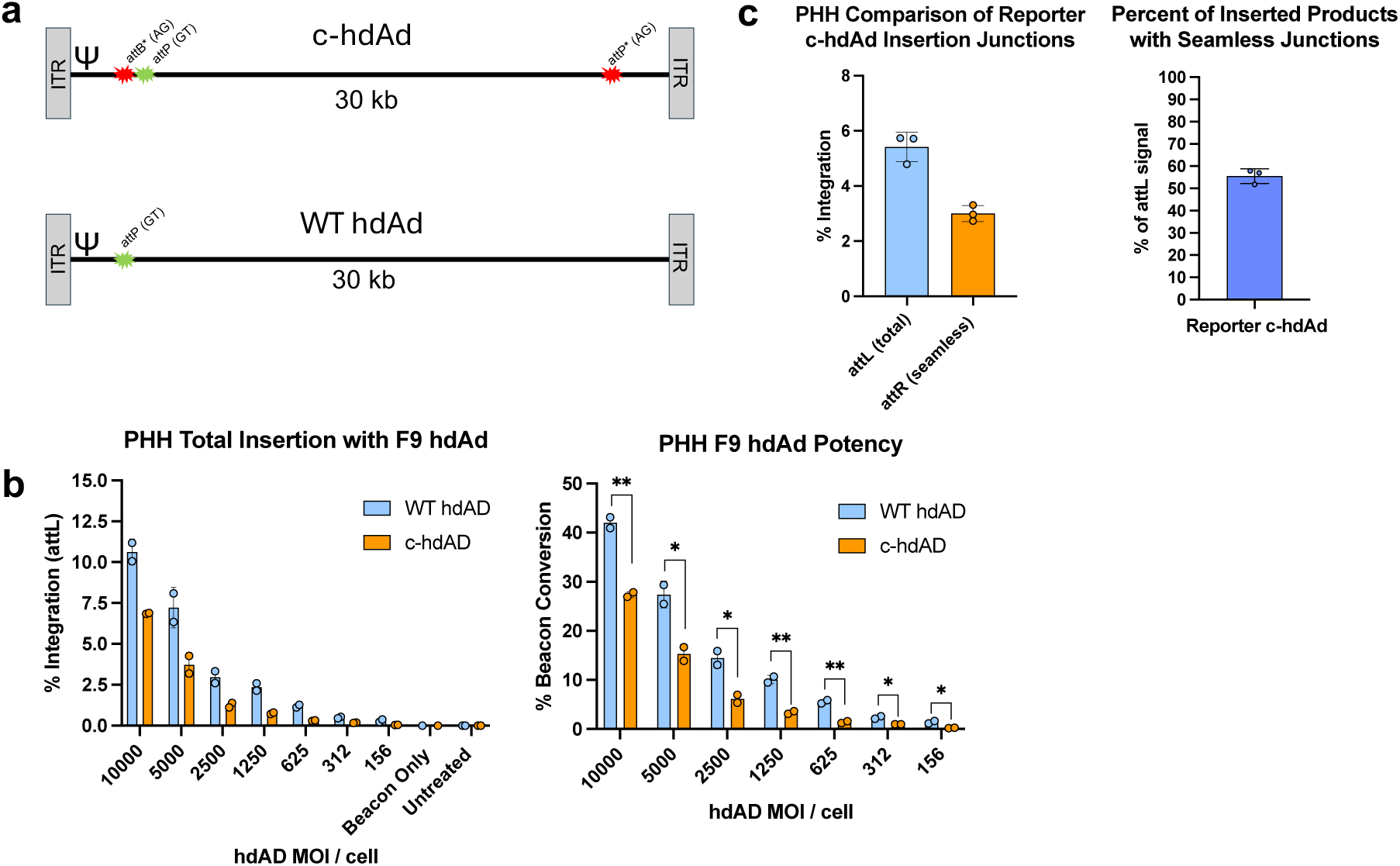
Validation of circle helper-dependent adenovirus in PHH. (**a**) Graphic depiction of circle hdAD (c-hdAD) and WT linear hdAD cargo. c-hdAD contains orthogonal attachment sites (AG dinucleotide) on the 5’ and 3’ ends of the genome. Bxb1 mediates intramolecular recombination to generate circular hdAD episome cargos. (**b**) Evaluation of F9 c-hdAd potency compared to WT F9 hdAd cargo at the *F9* locus in PHH. Statistical significance was calculated using Student’s unpaired two tailed t-test (* P<0.05, ** P<0.01). (**c**) Quantification of total and seamless insertion junctions following I-PGI with c-hdAD reporter cargo at the *F9* locus in PHH. Data reflects the mean and standard deviation of 2-3 biological replicates.

**Extended Data Figure 8.**
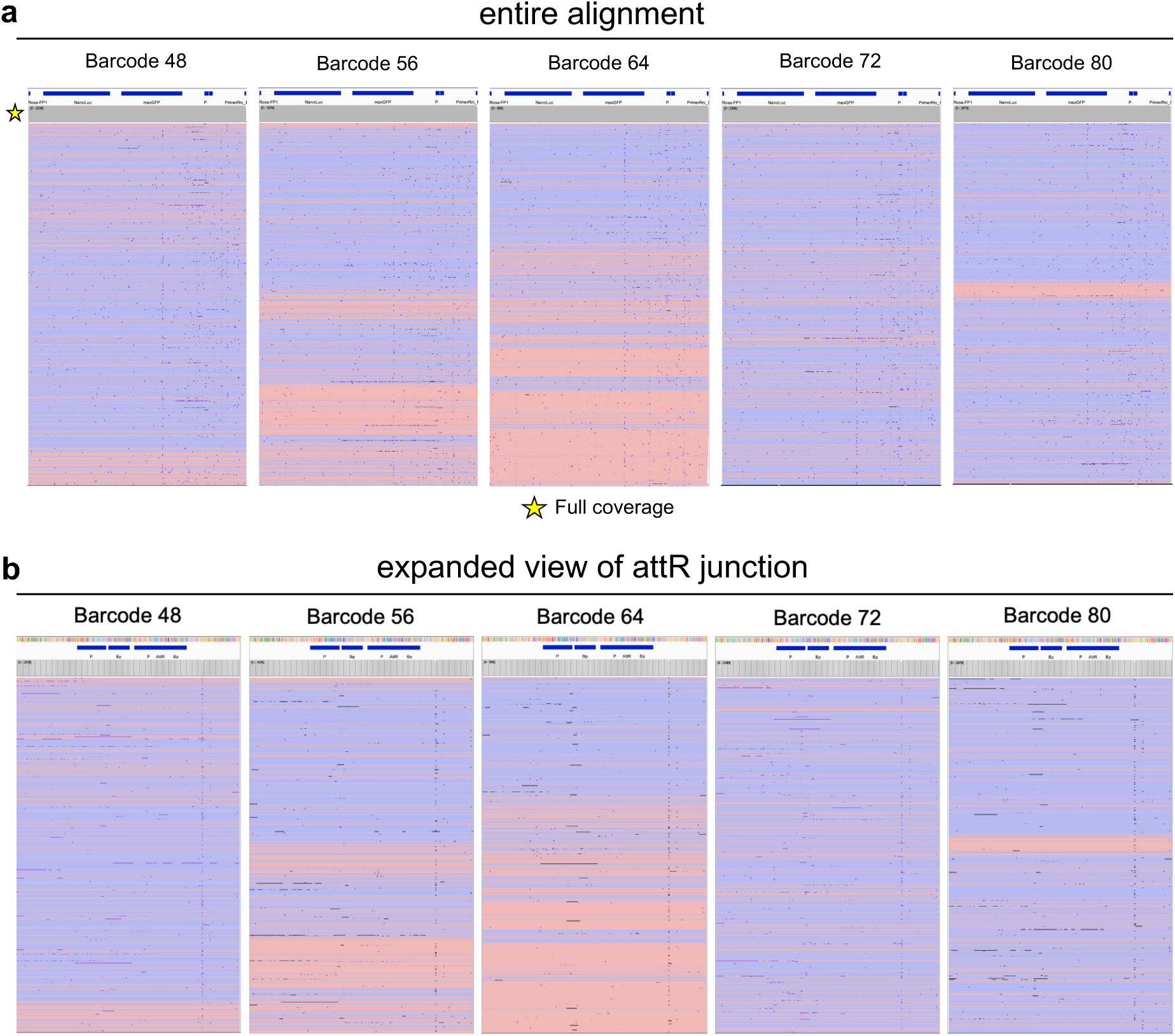
Sequencing of seamless cAAV recombination structures in mice. (**a**) Alignment of cAAV seamless insertion at the mouse *Rosa26* locus for each animal (barcode) in the cAAV group. (**b**) Expanded view of attR* / attR junction in the cAAV seamless insertion alignment for each animal. For a-b, each plot shows a collapsed read view where each line represents one read alignment in the forward (red) or reverse (blue) strand orientation.

**Extended Data Figure 9.**
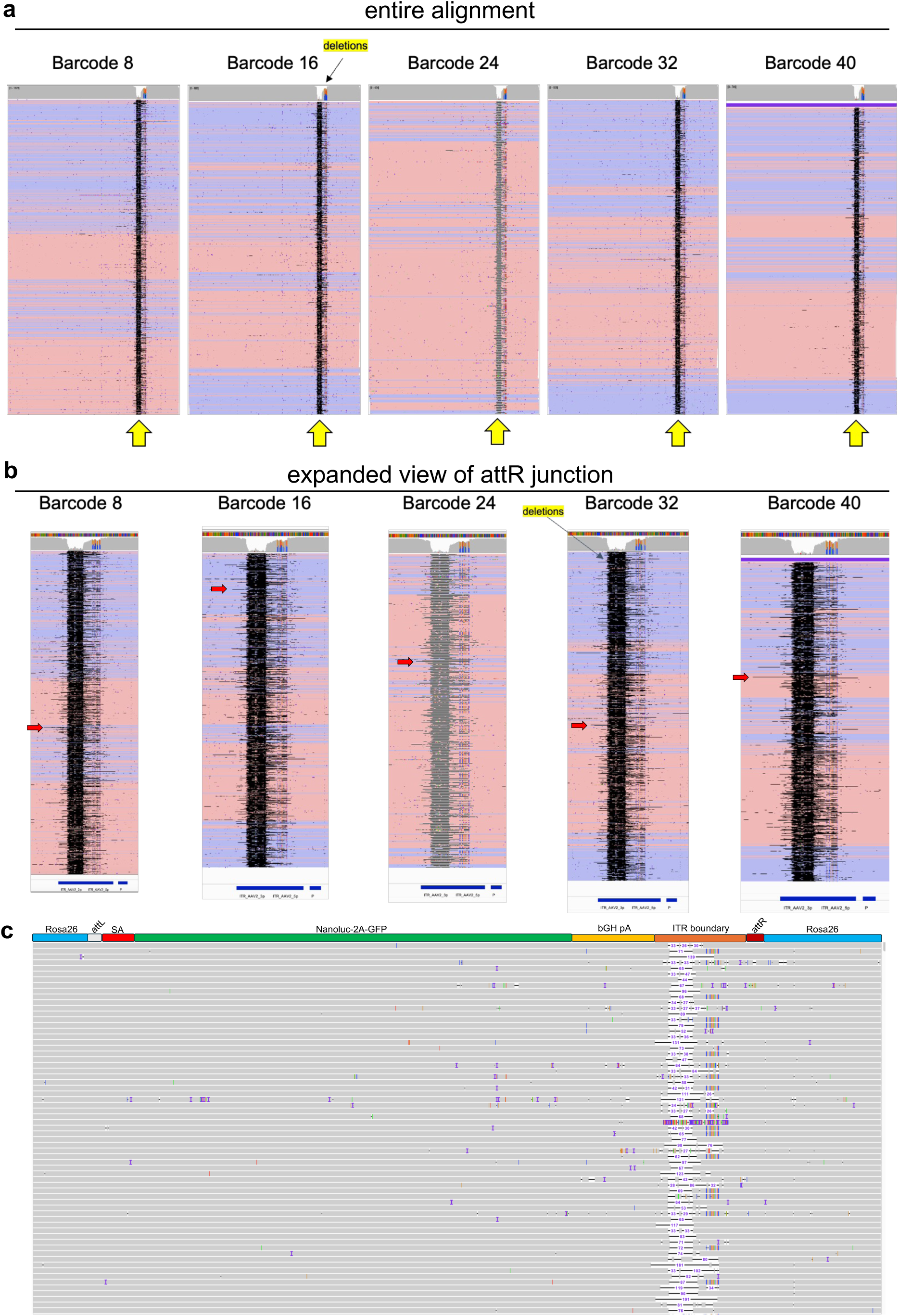
Sequencing of WT scAAV recombination structures in mice. **(a)** Alignment of WT scAAV insertion at the mouse *Rosa26* locus for each animal (barcode) in the control group. Alignment was performed using a full-length reference for linear scAAV. (**b**) Expanded view of attR* / attR junction in the linear scAAV insertion alignment for each animal. Deletions spanning into upstream cargo sequence are indicated with red arrows. (**c**) Expanded view of scAAV alignment in barcode 24. Each line represents an individual read with alignment (grey) and deletions (black line) of various sizes (purple number) within the ITR boundary region. For a-b, each plot shows a collapsed read view where each line represents one read alignment in the forward (red) or reverse (blue) strand orientation. _30_

**Extended Data Figure 10.**
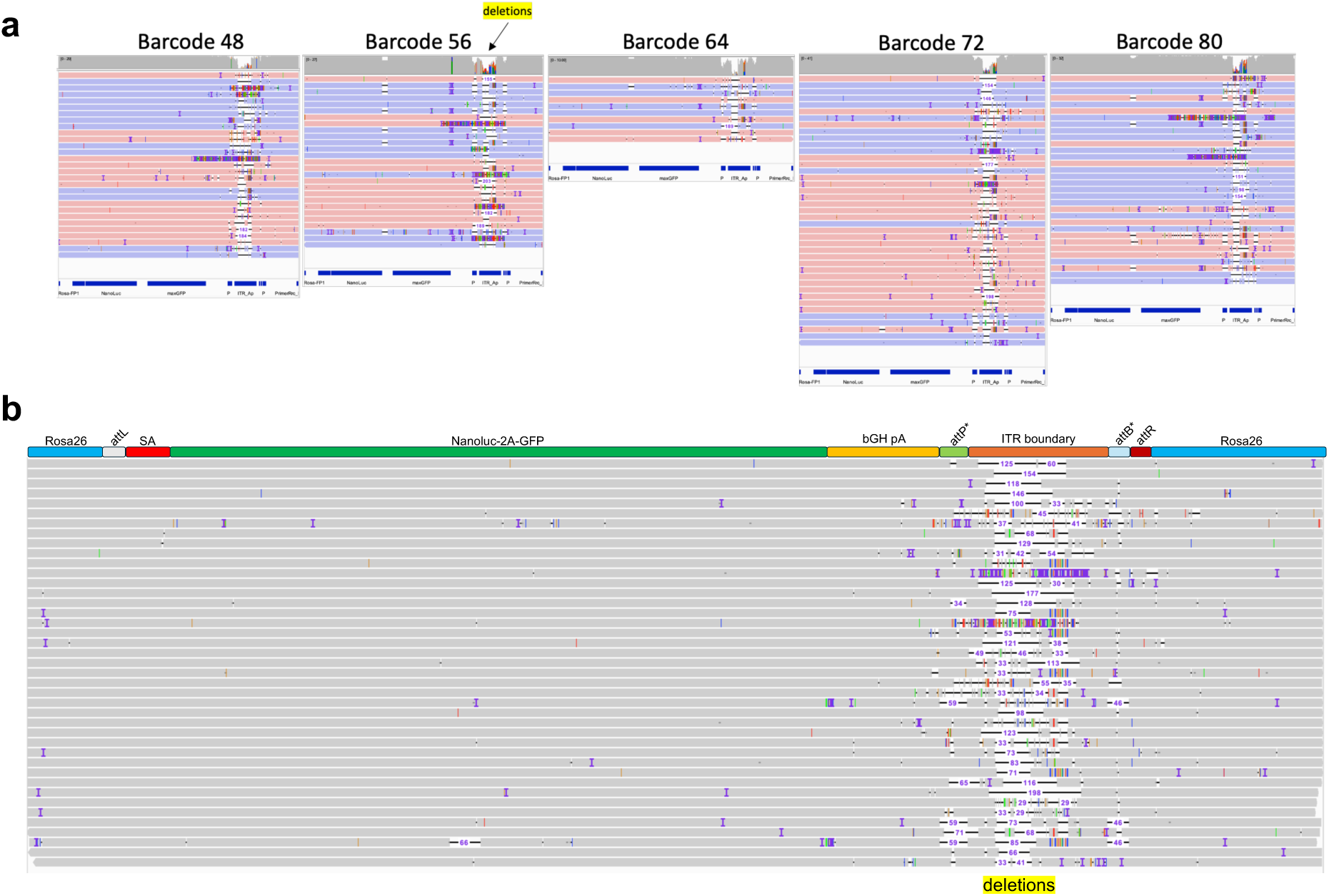
Sequencing of linear cAAV recombination structures in mice. **(a)** Alignment of linear cAAV insertion at the mouse *Rosa26* locus for each animal (barcode) in the cAAV group. Alignment was performed using a full-length reference assay for linear cAAV including ITRs. Each line represents one read alignment in the forward (red) or reverse (blue) strand orientation with deletions (black line) of varying lengths. (**b**) Expanded view of barcode 72 alignment where each line represents alignment (grey) containing deletions (black line) at varying lengths (purple number).

**Extended Data Figure 11.**
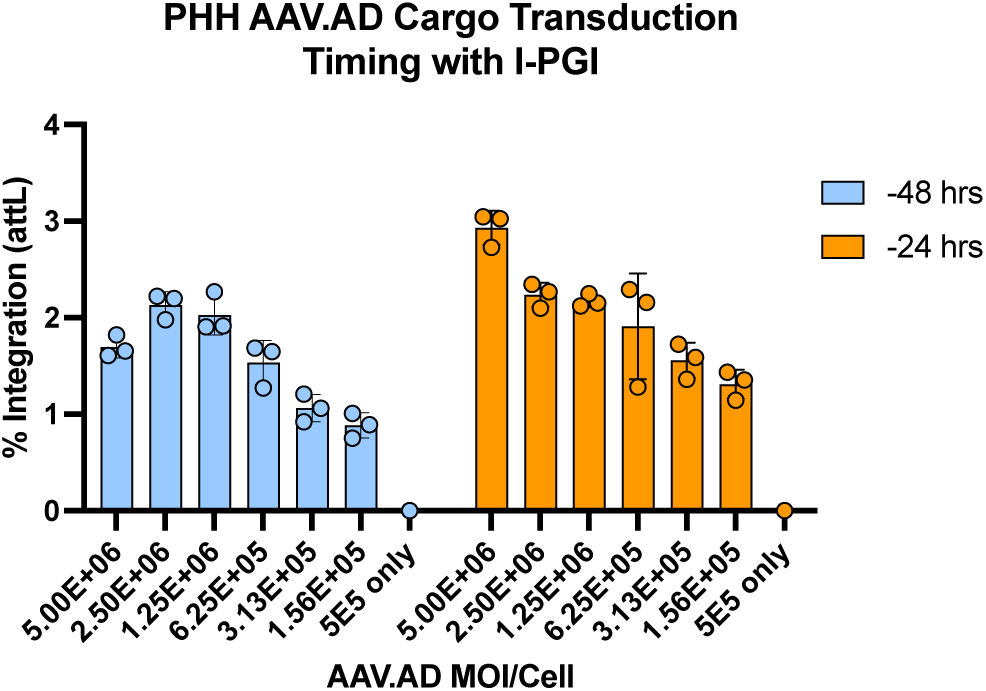
Extended predose timing of AAV.AD for I-PGI in PHH. The effect of modulating AAV.AD transduction timing on I-PGI outcome at the *PAH* locus. Transduction timing represents hours prior to single dose I-PGI transfection of synthetic atgRNAs, nCas9-RT and Bxb1 mRNAs. Data reflects the mean and standard deviation of 3 biological replicates.

**Extended Data Figure 12.**
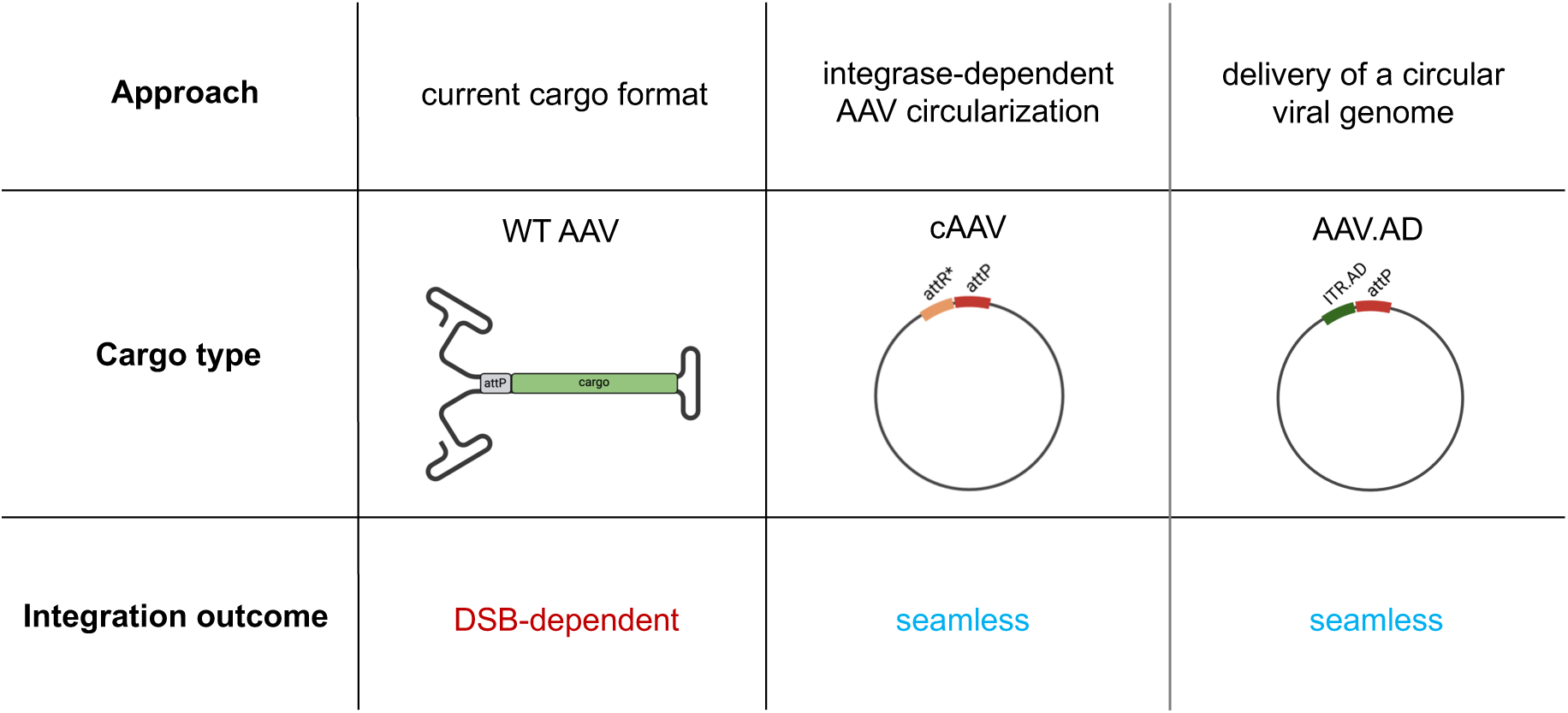
Summary of new circular AAV cargos and integrase-mediated insertion outcomes. Integration with wildtype (WT) linear AAV creates ITR-containing free ends that are likely recognized as a DNA double-stranded break (DSB) leading to DNA repair dependency. Circle-AAV (cAAV) utilizes integrase to circularize cargo and facilitates seamless DSB-free insertion. AAV.AD is a native circular genome that enables seamless DSB-free insertion.

